# A community machine learning challenge to predict the effects of gene perturbations on T cell differentiation for cancer immunotherapy

**DOI:** 10.64898/2026.05.21.726863

**Authors:** Jiaqi Zhang, Marc A Schwartz, Mohammed Mutaher, Oluwatomisin Olajide, Yuri Pritykin, Orr Ashenberg, Nir Hacohen, Caroline Uhler

## Abstract

Perturbations of genes with functional importance in T cells could be used to change the distribution of CD8 T cell states to enhance anti-tumor functions for cancer immunotherapies. We launched a world-wide computational challenge to predict the effects of gene perturbations and to devise objective functions for prioritizing gene perturbations that lead to desired T-cell state distributions. We supported the challenge by generating a single-cell Perturb-seq dataset profiling the effect of knocking out 73 individual expert-defined genes in T cells transferred into a mouse melanoma model. We compared the top algorithms developed by participants, and found that performance was primarily determined by the prior data used for gene feature representation, with perturbational data derived features, proving most effective. Experimental validation of the top 61 genes nominated by the algorithms revealed that perturbation of *Ndufv2* and *Dimt1* reached the defined objective and biased T cell differentiation toward desired states.

## Introduction

Decades of research into the molecular and cellular biology of T cells paved the way for today’s breakthroughs in cancer immunotherapy that have transformed patient care. The dominant treatment modalities, including immune-checkpoint inhibitors, adoptive cellular therapies, and cancer vaccines, broadly work to ensure that functional and active T cells infiltrate and clear the tumor.^1–4^ These immunotherapies have successfully targeted many tumor types, such as metastatic melanoma, renal cell carcinoma, non–small cell lung cancer, Hodgkin lymphoma, and B-cell leukaemias.^4^ Despite these remarkable successes, most patients fail to respond and many cancers remain refractory to these treatments, which has spurred research to understand and engineer more potent immune responses.

Within the tumor microenvironment, the T cell response spans a continuum of functional states that progressively shift from antigen-experienced progenitor states toward activation and then exhaustion/dysfunction states, ultimately leading to loss of tumor control by the T cells.^3,5^ The exhausted state is associated with poor immunotherapy outcomes and broadly defined by high expression of multiple inhibitory receptors (PD1, CTLA4, LAG3, TIM3), diminished cytotoxic activity, and disrupted cytokine signaling.

Immune-checkpoint blockade and adoptive cellular therapies are greatly affected by T cell exhaustion. Immune-checkpoint blockade enhances the cytotoxic response of tumor-specific T cells by reinvigorating the function of exhausted T cells and by recruiting new T cells to the tumor, while chimeric antigen receptor T cells (CAR T cells) are genetically engineered to recognize tumor-specific antigens and are being further engineered to avoid exhaustion through inactivation of PD-1 or TGF-β signaling.^3,5^ Furthermore the distribution of T cell functional states can affect the efficiency of cancer immunotherapies. For example, a strategy identified to improve the effectiveness of Immune Checkpoint Blockade therapy is to transiently increase the proportion of progenitor cells.^6^ For CAR T Cell therapy, reducing the proportion of terminal exhausted cells and increasing the proportion of progenitor, effector, and cycling cells has been found to improve the effectiveness of the therapy.^7^

Genetic perturbation could bias such proportions and enhance the anti-tumor potential of autologous T cells.^8,9^ Identifying the optimal genes to perturb remains a major challenge. The advent of Perturb-seq could provide opportunities to address this:^10^ Perturb-seq combines CRISPR/Cas9-based perturbations with a single-cell transcriptomic readout, allowing us to directly connect a genetic perturbation with its effect on a T cell’s functional state.^10,11^ However, it is unclear which are the most effective gene targets to include in a Perturb-seq experiment and testing all potential 20,000 gene targets in animal models or human systems is prohibitively expensive.

Machine learning (ML) models that predict gene perturbation effects in silico hold promise for experimental design, as they can help identify the most promising gene targets for further testing. However, developing reliable models is challenging, and there has been significant discussion in the literature regarding the best modeling approaches for Perturb-seq data.^12,13^ These debates highlight the difficulty of the prediction task and underscore the need for formal ML competitions to objectively benchmark the field’s ability to predict perturbation effects for meaningful downstream biological applications.

In 2023, we launched a global ML competition to foster the development of methods to predict the effects of gene perturbations on T-cell state proportions^14,15^. To support the challenge, we generated an in-house single-cell Perturb-seq dataset for the training, consisting of 71,393 cells measuring the transcriptional effects of 73 gene knockouts in naïve T cells following their injection into a mouse melanoma model. Here, we compare the performance of the top 20 algorithms and show the results of the experimental validation of the top genes predicted for each of the two objectives in the challenge. Our analyses revealed that the top-performing methods adopted a similar two-step mapping approach. Performance differences were mainly determined by the type of data used for gene feature representation, with features based on perturbational data (such as genome-wide Perturb-seq data on K562 cells and our own proposed T-cell specific perturbation features) outperforming those based on observational data. Experimental validation showed that two genes proposed by the algorithms, *Ndufv2* and *Dimt1* met the objectives of the challenge and significantly biased T cell differentiation toward a quiescent, progenitor-like state, yielding potential benefits for cancer immunotherapies.

### In-vivo single-cell CRISPR screens of genes affecting intratumoral T cell differentiation

Naïve T cells are activated in lymph nodes, enter the blood, and then infiltrate tumors where they differentiate into diverse T cell states, including a progenitor-like state that retains self-renewal capacity, an effector-like state with transient functionality, and a terminally exhausted state that is characterized by irreversible dysfunction (Figure 1A). We adopted a previously described *in vivo* CRISPR framework^16^ to knockout each of 73 genes in naïve CD8^+^ T cells, followed by transfer of perturbed cells into mice. 70 out of the 73 target genes were curated to include transcription factors known to be differentially expressed in T cells infiltrating melanoma samples^17–19^ and genes with known T cell regulatory functions, such as *Tcf7*, and their potential modulators^20^ (Table S1). The remaining 3 targets are known essential genes that serve as positive controls. We restricted the CD8^+^ T cell repertoire using OT-1 TCR transgenic mice, in which CD8^+^ T cells express a TCR specific for the ovalbumin-derived peptide OVA_257-264_ (SIINFEKL) presented by H-2k^b^, enabling recognition of B16-OVA melanoma cells that express ovalbumin. Naïve Cas9-expressing OT1^+^ CD45.2^+^ CD8^+^ T cells were transduced with a library of 257 single guide RNAs (sgRNA), including 3 sgRNAs per target gene for the 70 curated targets, 17 sgRNAs targeting 3 essential genes, as well as 15 non-targeting sgRNAs and, 15 intergenic-targeting sgRNAs that served as control.

**Figure 1.**
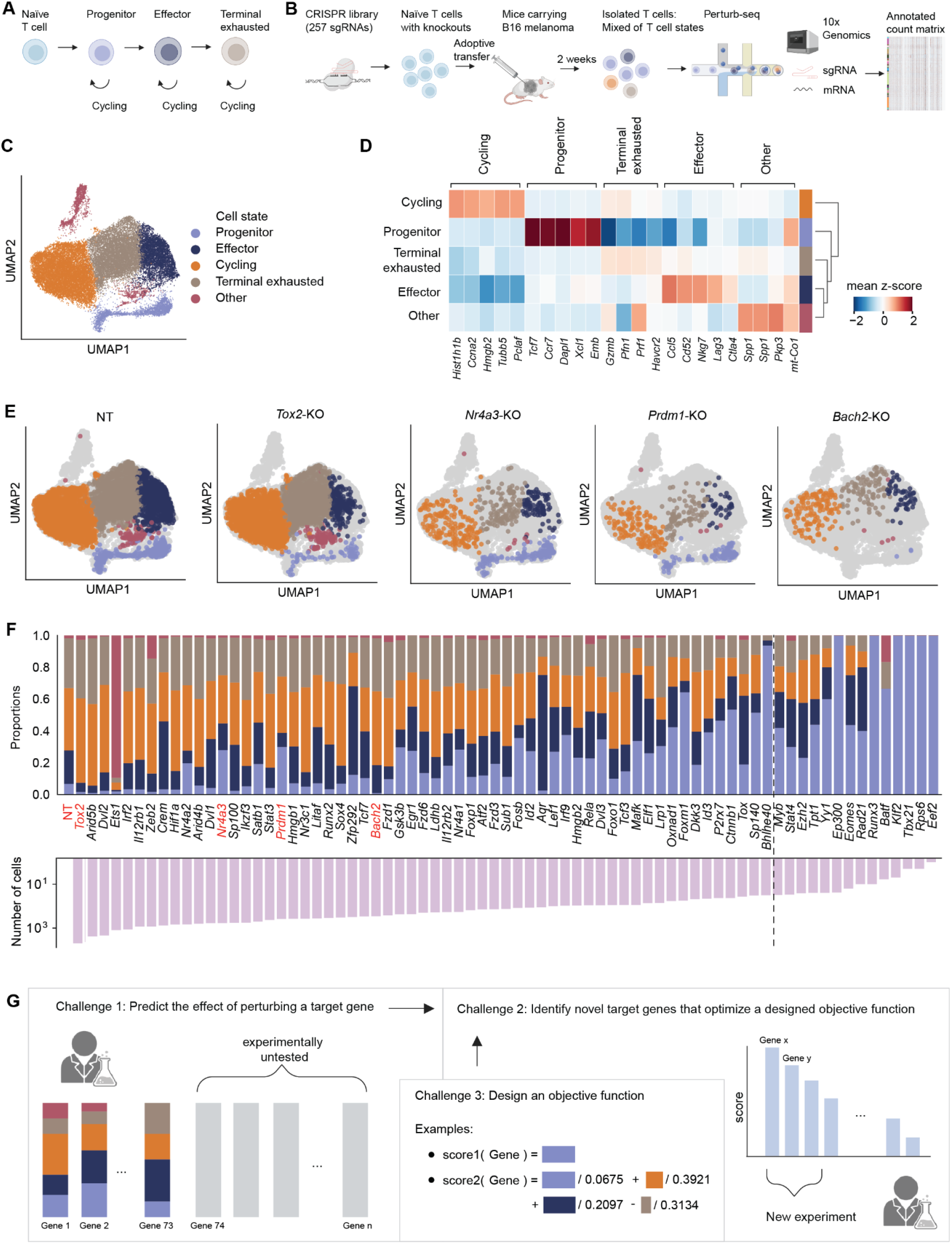
Overview of the cancer immunotherapy machine learning competition. **(A)**, Schematic representation of tumor-associated T cell states. The chronic exposure to weakly immunogenic tumor-associated self-antigens, together with other immunosuppressive mechanisms drives T cells into a hyporesponsive “exhausted state” with limited cytotoxic activity. **(B)**, Experimental design for the Perturb-seq experiment. Naïve Cas9 x OT1 CD45.2 CD8 T cells were transduced with a library of lentiviral vectors carrying single guide RNAs (sgRNAs) targeting 73 selected genes, and injected after 5 days into CD45.1 recipient mice, followed by implantation of melanoma. Tumors were harvested after 14 days and single-cell RNA sequencing was performed on the isolated CD45.2+ tumor-infiltrating Thy1.1+ perturbed T cells. **(C)**, UMAP visualization of the whole dataset colored by cell state. **(D)**, Heatmap showing the mean z-score for the top 5 marker genes used for the annotation of the 5 states. **(E)**, UMAP visualizations to highlight the cell state distribution for cell transduced with non-targeting sgRNAs (NT, on the left) or sgRNAs targeting genes with known functions (*Tox2, Nr4a3, Prdm1, Bach2*, from left to right). **(F)**, Bar plots indicating the proportion of cells in each state for the control (NT) and each single-gene perturbation (top) and the number of cells each perturbation has (bottom), with target genes shown in **(E)** highlighted in red. The cut-off of cell numbers below which we do not include as perturbations considered in the benchmark is indicated by the dashed line. **(G)**, Overview of challenge tasks. The participants are asked to predict perturbation effects on cell states of target genes that were held out from our dataset (Challenge 1), nominate untested perturbations predicted to induce desired cell state proportions (Challenge 2), and propose a quantitative objective function for ranking perturbations with respect to their effectiveness in inducing desired cell state proportions (Challenge 3).

After five days, the modified naïve T cells were adoptively transferred to congenic CD45.1^+^ mice, which were then subcutaneously injected with B16-OVA melanoma cells 1 day after T cell transfer to examine how the individual loss of genes would affect T cells in the tumor microenvironment (Methods). The tumor-infiltrating perturbed T cells were then collected after 14 days (when exhausted T cells accumulate) and analyzed by single-cell RNA sequencing (scRNAseq) (Figure 1B, Methods). We detected sgRNAs in 88.7% of the cells, and 49.0% of those contained only one dominant sgRNA based on counts, which we consider as cells with a single-gene perturbation. Among the 31,009 cells with single gene perturbations, we aggregated cells with guides corresponding to the same gene. By comparing transcriptional profiles of 15,077 measured genes across all perturbed genes, we identified three meta programs with convergent perturbational effects (Figure S1A). These gene programs were characterized by distinct molecular signatures and were enriched in pathways^21^ (Figure S1B) associated with T cell differentiation (meta program 1), proliferation (meta program 2) or activation (meta program 3).

Leiden clustering of the scRNAseq data identified five populations (Methods). We annotated four T cell populations as progenitor, effector, terminal exhausted, and cycling (Figure 1C), based on the expression of known gene markers, including: *Tcf7* and *Ccr7* for progenitors, *Ccl5* for effectors, and *Havcr2 (TIM-3)* for terminal exhausted T cells (Figure 1D, Figure S1C-D). The fifth cluster, annotated as “other”, included predominantly perturbation-specific cells, primarily from *Ets1*-knockout, along with cells that did not show significant expression of distinct cell-state-related markers (Figure S1E).

We confirmed that specific genes induced the expected changes in T cell proportions upon perturbation (Figure 1E,F). For instance, the depletion of *Bach2*, which enforces stemness and represses effector programs, reduced the proportion of progenitors.^22,23^ On the contrary, knocking out genes that promote activation, *Prdm1* and *Nr4a3*,^24^ led to an increase in the proportion of progenitor cells. Perturbation of essential gene positive controls, *Rps6* and *Eef2*, led to cell depletion (Figure 1F), while *Aqr* perturbation showed minimal depletion, suggesting that its essentiality does not extend to T cells.

Overall, the resulting distribution of cell states shows distinct proportions within different perturbations as compared to the non-targeting control (*NT*, Figure 1F). The dissimilarity in cell state proportions between perturbations targeting different genes is significantly greater than that observed between different guides targeting the same gene (*p*_*val*_ < 1*e* − 4, Figure S1F-G). Furthermore, we observed a correlation between the number of cells per perturbation and the proportion of certain cell states: negative for progenitor states (*r* =− 0. 83, *p*_*val*_ < 1*e* − 19, Figure S1H) and positive for cycling and terminally exhausted states (*r* = 0. 78, 0. 73, *p*_*val*_ < 1*e* − 15, 1*e* − 13, Figure S1H). This may be due to the intra-tumor progenitor cells being in a more quiescent, stem-like state with less proliferation than effector cells.

### Machine learning competition to accelerate algorithm development for cancer immunotherapy

Building on the perturbation dataset generated in-house, we launched the Cancer Immunotherapy Machine Learning Competition. This initiative aimed to unite teams of researchers from academia, industry, and non-profit organizations with the goal of developing algorithms to predict gene targets that could be manipulated to achieve specific T-cell state proportions more favorable for cancer immunotherapies.

We set up a three-part challenge (Figure 1G): **Challenge 1** – Build models based on the initial dataset to predict the cell state proportions for held-out gene perturbations. **Challenge 2** – Nominate novel, untested perturbations predicted to maximize an objective function described next in Challenge 3; these nominated perturbations were then evaluated in a follow-up experiment. **Challenge 3** – Design a quantitative objective function based on the predicted cell state proportions to rank perturbations by their effectiveness in inducing desired cell state compositions.

In response to **Challenge 1**, participants developed algorithms to predict cell state proportions caused by individual perturbations of seven randomly selected genes that were tested in our experiment but held out from the provided training datasets (Methods). The performance of the methods was evaluated based on the averaged absolute distance between the predicted cell state proportions and the available ground truth data. In **Challenge 2**, participants predicted the expected cell state proportions for the knockouts of all 15,077 expressed genes (Methods). In addition, they submitted two lists of genes ranked based on the predicted probability to score high in two objectives when perturbed. The first objective function was designed to capture a transient increase in the proportion of progenitor cells, reported to be beneficial for immune checkpoint blockade (ICB) therapy;^6^ the second was designed to capture a reduced proportion of terminal exhausted cells and increase in progenitor, effector, and cycling cells, reported to improve the effectiveness of CAR T-cell therapy.^7^ In **Challenge 3**, we asked participants to propose a third objective function potentially applicable also to other therapeutic strategies, aiming to identify more creative metrics for benchmarking ML predictions in a biological context (Methods). The proposed objective functions were evaluated through a single-blind peer-review process in which each submission was independently evaluated by three reviewers, followed by expert evaluations.

Most existing ML benchmarks are similar to the setup of Challenge 1, where evaluations are based on a held-out set of perturbations^25^. Here to test the ML methods for their application for biological efficacy, we selected the highest-ranked gene perturbations from the winning methods in Challenge 1 for experimental evaluation in Challenge 2. The experimental results were used to evaluate and rank all Challenge 2 submissions. Each submission from Challenge 2 was evaluated by ranking experimental-evaluated genes using the top objective function proposed in Challenge 3, in addition to the two ranked lists based on our example objectives.

Final winners for each challenge were selected based on the evaluation scheme described above. Out of the 246 total team submissions, we awarded prizes to the top 10 teams from Challenges 1 and 2, and the top 9 teams from Challenge 3 (Table S2). Overall, the winning methods of Challenges 1 and 2 used data from our Perturb-seq dataset, as well as various public databases, including STRING, NCBI, gene ontology, and other genome-wide Perturb-seq datasets for the training.^26–31^ These methods employed a range of ML approaches, including regression algorithms such as k-nearest neighbors, ensemble methods such as bagging, and deep learning methods such as generative models.^32–34^ The winning method of Challenge 3 proposed to incorporate uncertainty in estimating cell state proportions directly into the objective function via an upper-confidence bound strategy.^35^

### Benchmarking top participating methods for predicting cell state proportions

The top 20 methods, as ranked by the final results of both Challenge 1 and 2, were selected for further benchmarking and analysis (Methods). Fourteen out of these top 20 methods can be summarized by a two-step mapping, where the methods first learn gene feature representations of both tested and untested genes, then learn a map from gene features to cell proportions from the available data, and finally apply this to the gene features of the experimentally untested genes (Figure 2A). The other 6 methods either are very similar to the fourteen methods or cannot be clearly separated into distinct steps, but they perform worse than the 16 approaches that can be described by the two-step mapping in our benchmark (Figure S2A, Table S2). We discuss below these 14 two-step methods.

**Figure 2.**
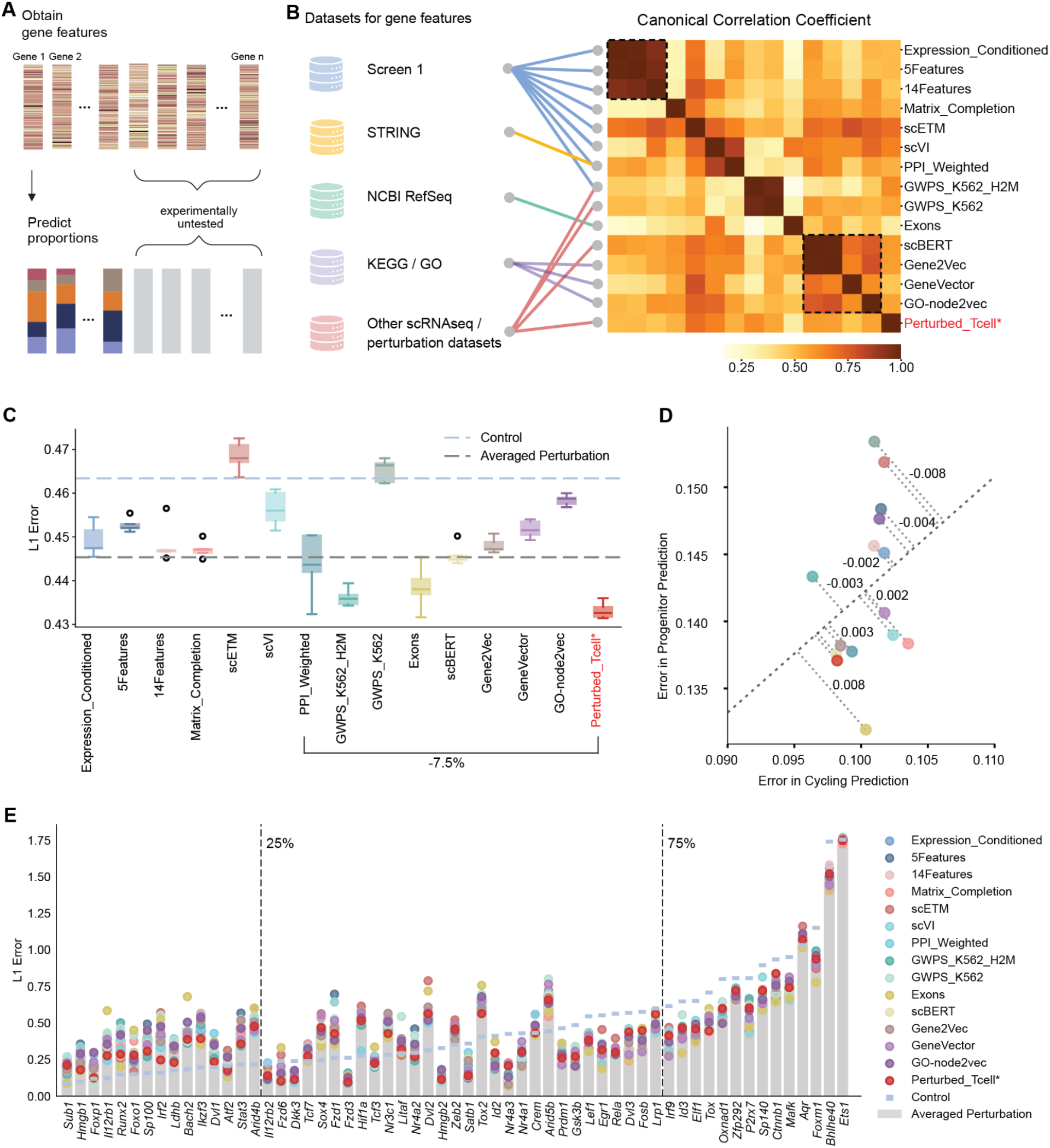
Benchmark of methods to predict gene perturbation effects. **(A)**, Schematic representation of how different methods predict cell state proportion vectors of experimentally untested genes. Each method obtains gene features for both experimentally tested and untested genes, then a map from gene feature to proportions is learned from the available data, finally it is applied to the gene features of the experimentally untested genes. **(B)**, Overview of gene features used by the methods in the benchmark. The left panel shows the source datasets from where the gene features are obtained. The middle panel links each individual gene feature to its source datasets. The right panel shows the similarity, measured by canonical correlation coefficient, of different gene features of genes from data, marked with two meta groups consisting of features derived from the provided Perturb-seq dataset and gene ontology databases respectively. **(C)**, Boxplot showing L1 errors calculated on a cross-fold split, repeated five times, by combining a nearest neighbor predictor with each of the 15 features used by the top performing methods indicated on the X-axis, plus our proposed feature gathered from perturbation datasets on T cells in the literature (Perturbed_Tcell, in red). The origin of the acronyms and their corresponding submissions are given in Table S2. Prediction using the control and the averaged perturbation baselines are shown in the dashed line. **(D)**, Comparison between error in progenitor proportion prediction versus cycling proportion prediction of different methods. The corresponding method of each colored dot follows the same legend in **(E)**. The dashed line is obtained by linear regression between two errors and the annotations are the distances between the dots and their projection on the dashed line. **(E)**, Breakdown of the averaged *l*_1_ errors on different perturbations. Each method’s performance on each perturbation is indicated by a dot, where the averaged perturbation baseline is indicated by the grey bar.

For the first step, features were extracted based on multiple resources, derived from either the provided Perturb-seq dataset or online databases such as STRING, NCBI, gene ontology, and other genome-wide Perturb-seq datasets^26–31^ (Figure 2B). To assess the similarity of the proposed features, we compared them using canonical correlation analysis, which for each pair of feature vectors finds the directions that best correlate this pair after projection (Methods). These 14 features show several meta groups (Figure 2B). Many features from the same meta group are derived from the same database, which may explain the observed correlations (Figure 2B). For example, one meta group consists of features derived from the provided Perturb-seq dataset, while another meta group contains features derived from gene ontology databases. We also proposed an additional curated feature, the Perturb_Tcell feature, obtained by integrating prior perturbation data on T cells that we sourced from the literature^36–44^ and selected based on the provided Perturb-seq data in a cross-validation setup (Figure 2B, Methods).

For the second step, several predictors can be used, which fall into two categories: (1) direct prediction of cell state proportion, which maps the feature vector directly to the T-cell state proportion vector e.g., using nearest neighbor regression; (2) transcriptomic-based prediction, which first maps the feature into a distribution of predicted transcriptomic measurements at single-cell level e.g., using a generative model, then classifies each generated single cell into five T cell states using the transcriptomic measurements e.g., using MLPs. Transcriptomics-based prediction generally performs worse than direct prediction in our benchmark (Figure S2A). This suggests that direct prediction more effectively captures the target signal without going through an intermediate step where errors may compound and the target signal is diluted, a phenomenon characterized by the Data Processing Inequality.^45^ Therefore, we focus on direct prediction methods in the following analyses.

We repeated the train-test split six times, so that each perturbation is held-out once (Figure 2C, Figure S2B). For each split, we trained each predictor and selected its hyperparameters using a cross-validation scheme on the training set, and we then evaluated it on the perturbations in the test set (Methods). For each perturbation, we considered the following evaluation metrics: (1) average absolute loss (i.e., L1 error) across all T-cell states, and (2) absolute loss on key states such as progenitor and cycling (Figure 2C-D). We then averaged the L1 error across all perturbations to evaluate a method’s overall performance (Methods). For a robust comparison, we tested the performance of the same predictor, nearest neighbor regression, using the different features (Figure 2C-E). Alternative predictors, including kernel regression and bagging were tested too, showing similar results (Figure S2D). We included two baselines, one that simply predicts the proportion of T cell states in the non-targeting control cells, and one that predicts the averaged proportions of T cell states across all training perturbations, which has been shown in the literature^46^ to be competitive against many deep learning solutions.

While the gap between the best and worst performing features is small (7.5%), features based on exons (derived from the NCBI Gene Database),^47^ genome-wide Perturb-seq (GWPS),^48^ and our own curated feature, Perturbed-T-cell, outperform others; no other prediction method is significantly better than the averaged perturbation baseline (Figure 2C). Two of the best performant features, GWPS-based and our own curated Perturbed-T-cell based features, are derived from external perturbation datasets. We investigated these two features further (Methods). We found that the genes leading to similar or dissimilar effects on T cells tend to also lead to effects of the same nature on K562 cells (Figure S3A), suggesting why using perturbational data on K562 cells is helpful for prediction on T cells. The collection of previous perturbation data on T cells share information with shifts caused by perturbation in our data; differentially expressed signals from prior data are most relevant for shifts in earlier states like progenitor and effector (Figure S3B).

We also investigated the impact of the combined (Figure S3C-E), or individual exon features (Figure S4) on the predicted on-target effect size (log-fold change), and independent measures of gene activity and essentiality, using our dataset, GWPS, X-atlas, unperturbed PBMCs, and DepMap.^48–51^ Our analysis revealed that exon features are correlated with the on-target effect size across our dataset, GWPS, X-atlas, and DepMap (Figure S3C-E, Top Panel; Figure S4D-E). We also observed correlations between exon features and gene expression across these perturbational datasets, and on an unperturbed PBMCs dataset (Figure S3C-E, Bottom Panel; Figure S4C-F), a phenomenon recognized in the literature that exon number and size are negatively correlated with expression level.^52^ The correlations are mostly observed in exon count, mean length, and maximum length (Figure S4). As on-target effect size is often associated with gene expression, we were wondering if the observed correlation between these exon features and on-target effect size is mediated by gene expression. Therefore, we conducted conditional independence tests using our dataset. Interestingly, we found these exon features to be independent of on-target effect size given gene expression (*p*_*val*_ > 0. 05 for the null hypothesis of independence, Methods). This suggests that the correlation between exon features and on-target effect size can likely be explained by gene expression. Such correlation might enable exon features to better predict states like progenitor cells (which have fewer regulatory programs and lower transcriptional variability), while they may struggle with more complex states, such as cycling cells (Figure 2D).

To further understand how the different methods generalize to different perturbations, we assessed the performance of the methods by each perturbation (Figure 2E). We observed that the L1 error per gene is highly correlated with the L1 distance between the perturbed T cell distribution and the mean effect of the training set perturbations (i.e., the L1 error of the averaged perturbation baseline). This is expected, as the learned parameters of each method are inherently optimized to predict output that are similar to the training data. It is worth noting that prediction errors are largely driven by the perturbed data rather than the control population. For perturbations that resulted in minimal change compared to control (bottom 25% in terms of L1 distance to control), all prediction methods performed worse than the control baseline, as the control itself provides a strong estimate for these cases. In contrast, for perturbations that deviated substantially from control (top 25% in terms of L1 distance to control), most methods outperformed the control baseline by a large margin, since the averaged perturbation effect better captures their true response. For intermediate perturbations (in the 25%-75% range), performance depended on whether the averaged perturbation effect or the control was closer to the true perturbation profile. Notably, our constructed Perturbed_Tcell features consistently rank among the top-performing methods across perturbations.

### Prediction on a public T cell perturbation dataset

To further confirm the comparative results, we ran these methods on a public dataset.^53^ This dataset was collected using the same mouse model and a similar procedure, but (1) T cells were activated before being injected into the mice, and (2) T cells were harvested 7 days as opposed to 14 days after adoptive transfer. Therefore the focus was on genes that act relatively earlier. To keep the task comparable to that in our competition, we aligned the public dataset with our dataset using Harmony for batch correction^54^ (Methods, Figure 3A), and annotated it with our defined cell states by mapping each cell to the nearest state cluster (Figure 3B). The public dataset targeted 180 TFs chosen based on differential expression analysis on prior datasets. Among the 180 TFs, 36 TFs were also tested in our dataset. For the perturbational effects of these 36 TFs, we observed a consistent trend toward a higher proportion of progenitor cells in the public dataset compared to ours (*p*_*val*_ < 1*e* − 4, Figure S5A), which is apparent already when comparing the non-targeting control (0.406 versus 0.067), reflecting differences in experimental procedures. In addition, we also observed that the perturbation effects in terms of L1 distance between perturbed cell state proportion and control is significantly greater in our data versus the public dataset, indicating that on average we have larger perturbation effects (*p*_*val*_ < 1*e* − 4, Figure S5B).

**Figure 3.**
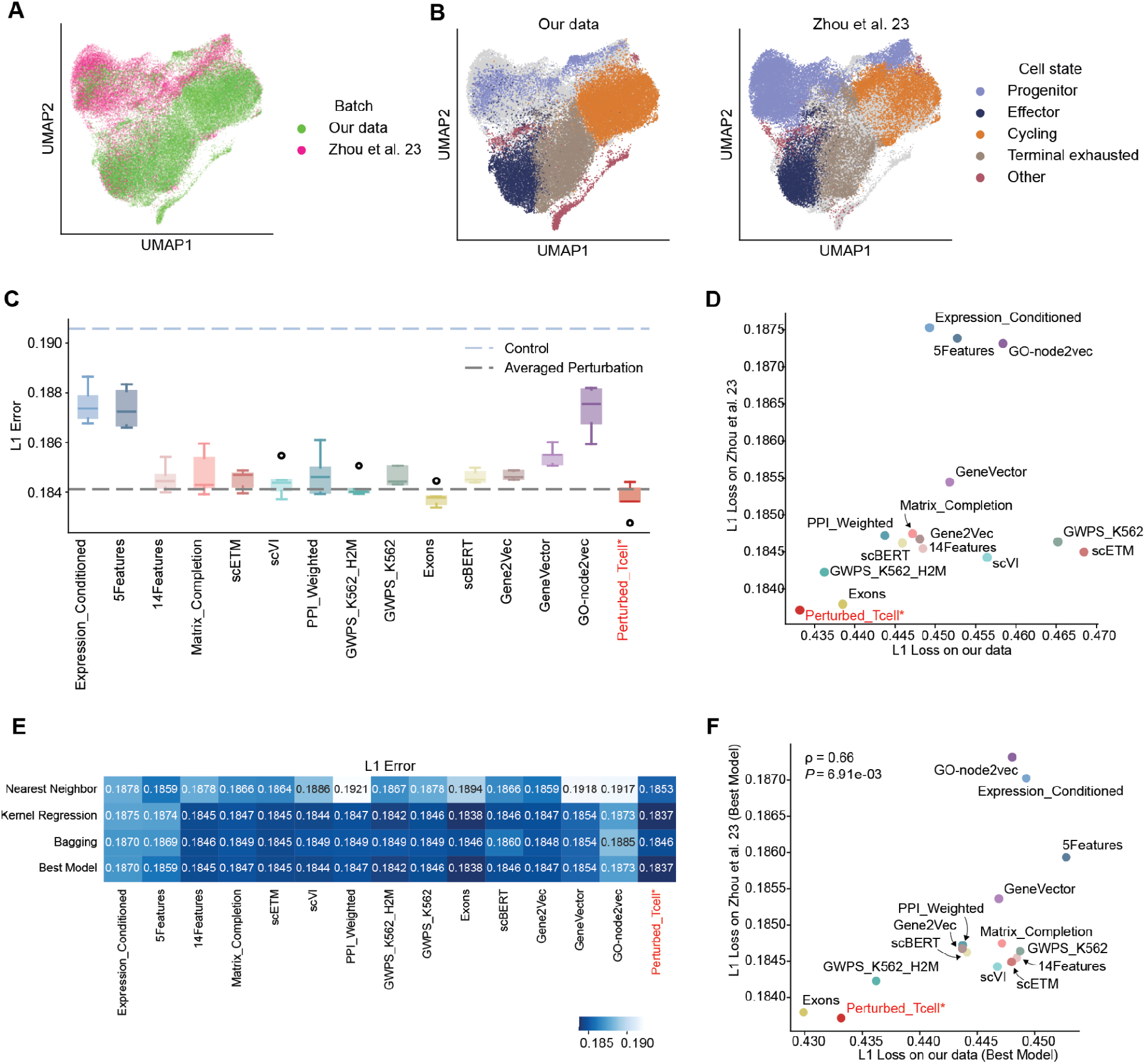
Benchmarking results on a public T-cell Perturb-seq data. **(A)**, UMAP visualization of our data and the public Perturb-seq data^53^ after batch correction. **(B)**, UMAP visualization highlighting the cell states in our data, and annotation based on our cell states in the public Perturb-seq data.^53^ **(C)**, Boxplot showing *l*_1_ errors of different features using the kernel predictor, across five repeated runs on a cross-fold split on the public Perturb-seq data.^53^ Prediction using the control and the averaged perturbation baselines are shown in the dashed line. **(D)**, Comparison of averaged *l*_1_ errors of different methods in our data from Figure 2C and the public Perturb-seq data^53^ in **(C)**. **(E)**, Averaged error of different methods, across combinations of gene features and predictors (first three rows). The last row shows the best predictor of each gene feature from the first three rows. **(F)**, Comparison of averaged *l*_1_ errors of different methods with the best predictor in our data and the public Perturb-seq data^53^ (Spearman ρ = 0. 66, *p*_*val*_ < 1*e* − 2).

We applied the methods obtained by combining each of the 15 features with the same predictor (nearest neighbor, kernel regression, or bagging) on this annotated dataset following the same train-test procedure and metrics described above. By comparing different features, using the kernel regression predictor that generally works better on this dataset, we confirmed a similar trend with several features, namely, our proposed Perturbed_Tcell feature, GWPS based, and exon based features, performed better than other features and the averaged perturbation baseline, while other features are not better than the averaged perturbation baseline (Figure 3C), possibly because the perturbation effects in this dataset are generally smaller than in our dataset (Figure S5B). By directly comparing the rank of different features on this dataset versus ours, we see that the ranks are mostly preserved if we use the kernel regression predictor (Figure 3D) and well preserved if we take the best predictor among the nearest neighbor, kernel regression, and bagging (ρ _*val*_ = 0. 66, *p* < 1*e* − 2, Figure 3E-F, Figure S5C).

### Identification and ranking of target genes towards desired states

Next, we ranked the genes submitted in Challenge 2, by the Challenge 1 winning teams (Table S2). For each team, based on the predicted cell state proportions for the remaining untested targets, we used two objective functions that take in a proportion vector and output a scalar score to rank the untested targets. The two objective functions are constructed to resemble potential desired cell states for ICB and CAR-T therapies.^6,7^ Namely, as illustrated for the tested genes using ground-truth data (Figure 4A-B), the ICB objective aims to transiently increase the proportion of progenitor cells while maintaining a certain population of cycling cells (Figure 4A, Methods). The CAR-T objective reduces the proportion of terminal exhausted cells and increases the proportion of progenitor, effector, and cycling cells (Figure 4B, Methods).

**Figure 4.**
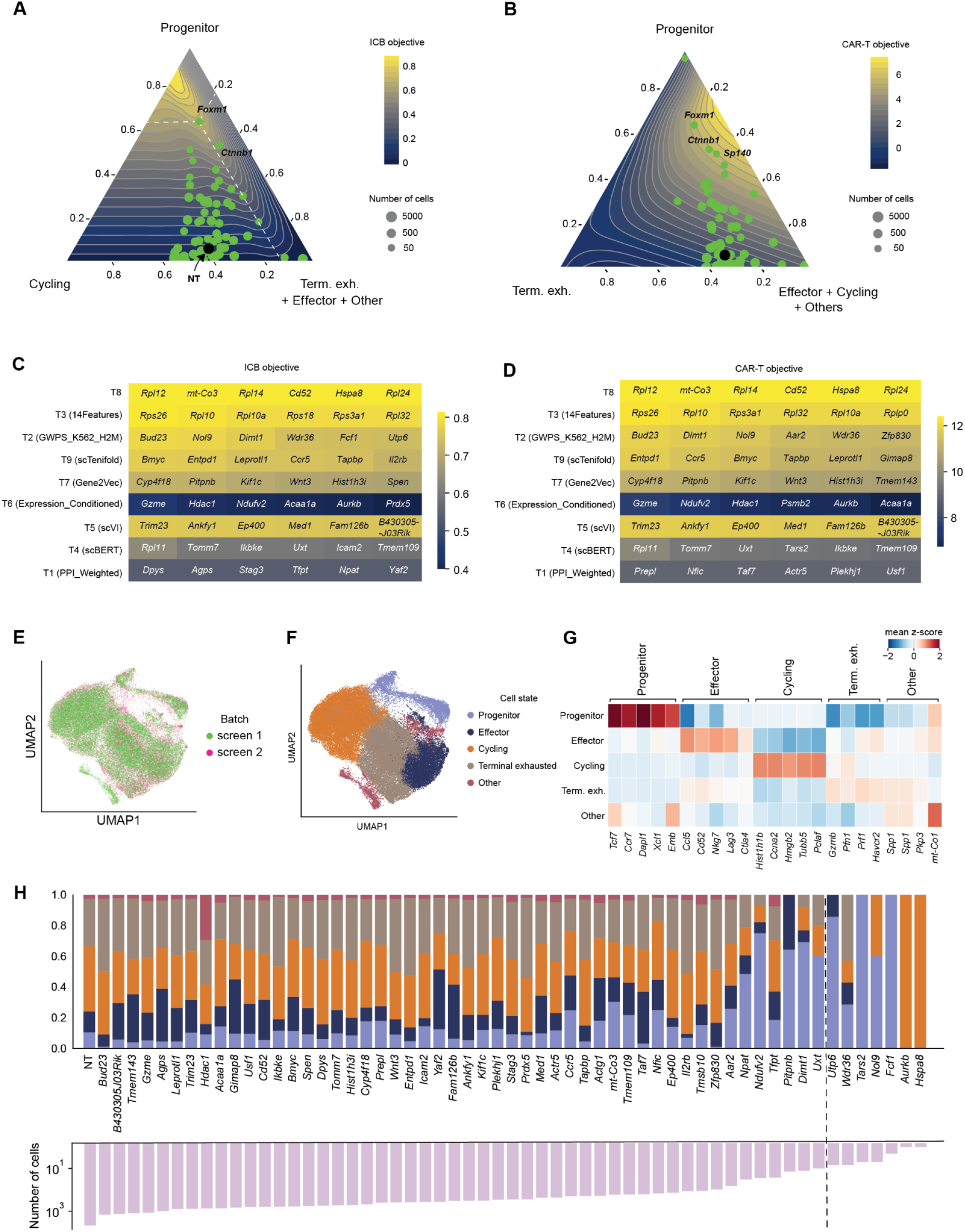
Objective functions and the validation screen. **(A-B)**, Ternary plot visualizing perturbations in our first screen in terms of the ICB objective in **(A)** and CAR-T objective in **(B)**. The heatmap coloring indicates the smoothed averaged objective function values of the proportions at the corresponding spot. Each dot corresponds to a perturbation (or NT, as annotated by the black dot), where the size of the dots indicates the number of cells for that perturbation and top 2 perturbations of each objective function are annotated. In **(A)**, perpendicular lines of *Foxm1* are illustrated to aid reading the axes (left axis corresponds to proportion of progenitor, bottom axis corresponds to proportion of cycling, right axis corresponds to the added proportion of the remaining states). **(C-D)**, Nominated genes for the follow-up experiments, which are top genes for the ICB objective in **(C)** and CAR-T objective in **(D)** predicted by the teams that are winners of challenge 1 and from which the benchmark methods are derived. Color indicates the predicted objective function values for each gene. **(E)**, UMAP visualization of the validation screen (screen 2) and the first screen (screen 1) after batch correction. **(F)**, UMAP visualization of both datasets colored by cell state. **(G)**, Heatmap showing the mean z-score on screen 2 for the top 5 marker genes of the 5 states from screen 1. **(H)**, Bar plots indicating the average proportion of cells in each state for the control (NT) and each single-gene perturbation (top) and the number of cells each perturbation has (bottom) in screen 2. The cut-off of cell numbers below which we do not include as perturbations considered in the benchmark is indicated by the dashed line.

In Challenge 3, we asked the participants to propose new objective functions by submitting report-style writeups and evaluated the submissions based on a peer review process (Methods, Table S2). The top submission proposed an objective function that incorporated an uncertainty term (a lower confidence bound, inspired by the bandit literature^35^), tied to the number of cells observed in each perturbation. Because the number of cells for untested genes was unknown a priori, this objective was not used to select genes for the validation screen. However, it was applied post-hoc to rank the validated target genes (using the observed cell counts), and this ranking was subsequently used, along with the rankings from the other two objectives, to score all Challenge 2 submissions (Table S2).

In summary, for the validation screen, we selected the top genes predicted by the methods developed from the Challenge 1 winning teams (the same from which we derived the benchmark methods) for the ICB and CAR-T objectives (Figure 4C-D, Figure S6A-C). We excluded several DepMap essential genes^50^ that are involved in proteasome and ribosomal families which are known to lead to cell death (Table S1). In total, we selected 57 target genes (the majority of these genes are not well-known T cell regulators) and designed 3 guides per gene (Table S1). We performed a Perturb-seq experiment of these target genes following the same protocol (Figure 1B) and the same preprocessing steps (Methods, Figure S6D) as in our first experiment. We then annotated the cell states by first batch correcting between the validation screen (screen 2) and our initial screen (screen 1) then assigning cells to the states of screen 1 based on majority vote of the neighboring cells (Figure 4E-F, Methods, Figure S6E). Overall in the second screen, we observed adequate coverage of the four cell states, namely progenitor, effector, cycling, and terminal exhausted (Figure 4F-G), but the cell state “other”, which was mainly driven by *Ets1*-knockout in screen 1 was less frequently observed in screen 2. The resulting proportions of cell states of different knockouts in screen 2 as well as the number of cells are shown in Figure 4H.

### Experimental validation of predicted top target genes

After quality control and data processing, we ranked the genes tested in our validation screen (screen 2) according to the three objective functions. Since the ICB objective has the same functional form for both screens, we can directly compare the genes tested in screen 2 with those tested in screen 1 (Figure 5A). We found that the expert-selected targets tested in screen 1 were generally superior and already covered high-fidelity targets. Nonetheless, the ML-guided approaches identified two targets *Dimt1, Ndufv2* with higher ICB objective, which led to less terminal exhaustion and more progenitor states than all targets tested in screen 1 (Figure 5B), showing that ML-guided approaches can identify rare targets that outperform current expert’s knowledge.

**Figure 5.**
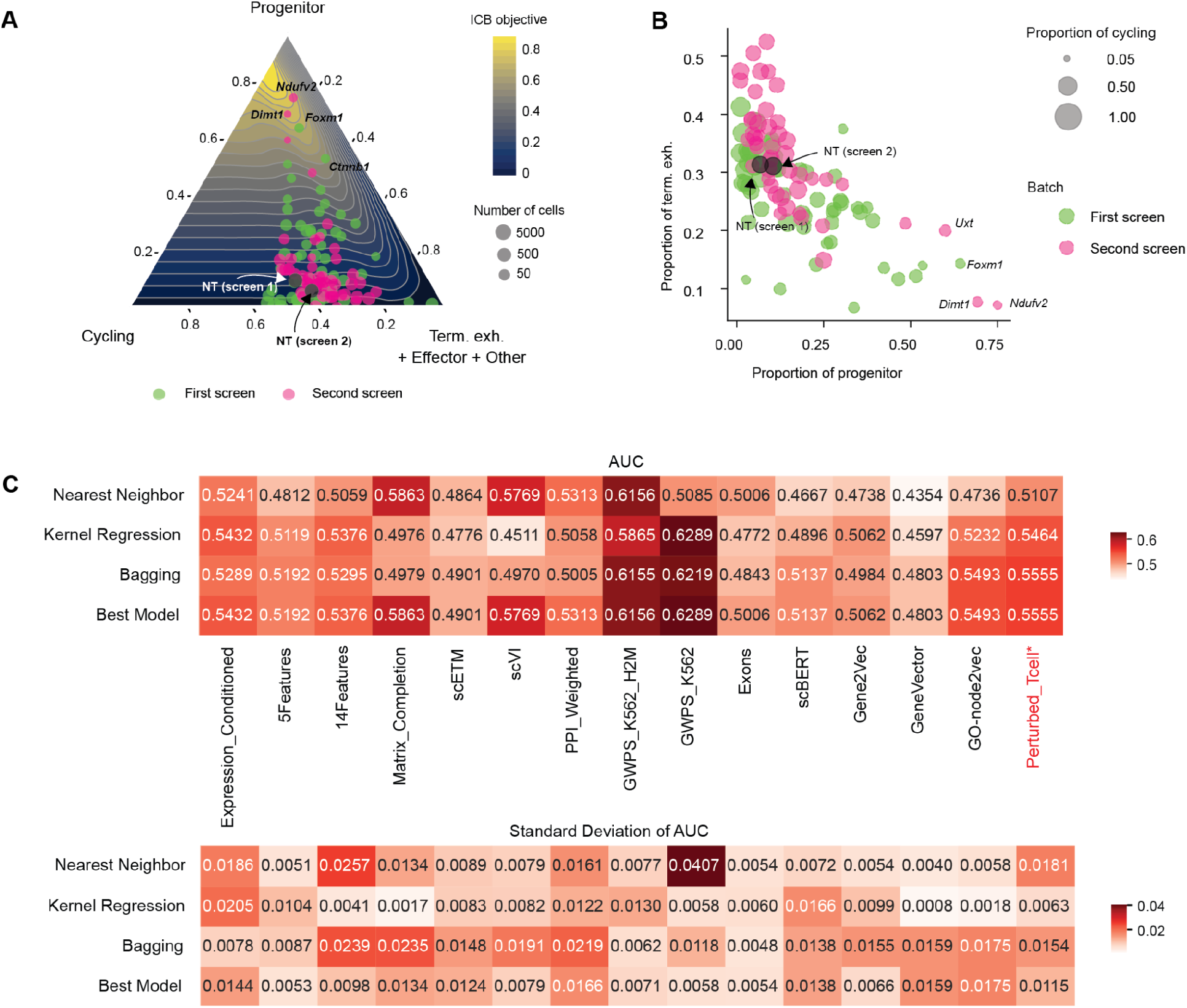
Comparison of both screens in terms of ICB objective and validation of methods on the screen 2. **(A)**, Ternary plot visualizing perturbations in both screens in terms of the ICB objective. The heatmap coloring indicates the smoothed averaged objective function values of the proportions at the corresponding spot. Each dot corresponds to a perturbation (or NT, as annotated by the black dot), where the coloring indicates the screen, the size of the dots indicates the number of cells for that perturbation and top 2 perturbations of both screens are annotated. **(B)**, Scatterplot of perturbations in both screens comparing proportions of progenitor against proportions of terminal exhausted, with the four perturbations annotated in **(A)** and NT controls annotated. The coloring indicates the screen, and the size of the dots indicates the proportion of cycling proportion. **(C)**, Validation of Methods when trained only on screen 1 to rank the perturbations in screen 2 according to the ICB objective. Top plot shows the area under the curve when comparing the predicted order to ground-truth order, averaged across 5 runs. Bottom plots show the standard deviation across 5 runs.

We further looked at whether the benchmarked prediction methods could perform well in a situation that mimics the exact procedure of the experiments, i.e., uses the data from screen 1 to rank the targeted genes in screen 2 for the ICB objective (Methods). We used the same metric (AUC for the two ranked lists, one predicted by the methods, one given by the ground-truth data from screen 2) to re-evaluate the different models (Figure 5C, Methods). The top performing methods were those using the GWPS-derived features, suggesting that the gene features derived from this perturbation screen contained the most useful information to rank the validated gene set, despite the screen being performed in a very different environment (in-vitro K562 cell line as opposed to T cells infiltrating melanoma *in vivo*), and on a different species (human as opposed to mouse). The other two top performing features, scVI and matrix completion, both used all perturbation data from screen 1 to derive features. The top feature in the previous benchmarks, Perturbed_Tcells, which was proposed by us, was still amongst the top performing methods but was not dominating anymore, potentially because the previous perturbation screens primarily targeted the space of known regulators but was relatively less informative for unknown genes that were proposed by the ML methods tested in the validation screen, while the other priors that rely less on perturbational information contains equal information for known regulators and unknown genes. The exon features, which we discovered primarily reflect expression-mediation, were not performing well in this task and had a performance closer to random guessing (0.5).

Since the evaluation procedure was the same as in the competition except with the addition of the 7 perturbations that were held out as test set in Challenge 1, we could directly compare the re-implemented methods in the benchmark to the officially submitted version. In most cases, the re-implemented best model works better than the submitted version with the additional data, and the ordering between most of the reimplemented versions is consistent with the submitted version, with the exception of a few methods, including GWPS_K562, scVI, and Matrix_Completion, which show a large variance when different prediction algorithms were used (Table S2, Figure 5C).

WWith respect to the CAR-T objective, we found the same two top target genes (*Dimt1, Ndufv2*) and a similar trend in terms of the methods’ performance (Figure S6F-G). For the participant-proposed winning objective, its ranking of target genes can be found in Table S3. We however did not predict it using just screen 1, since it requires predicting the cell numbers.

### *Ndufv2* and *Dimt1* shift the differentiation towards more progenitors

For the two hits, we sought to understand why they led to higher objective function values. From the control cell population in screen 1, we saw that both *Ndufv2* and *Dimt1* are significantly highly expressed in cycling cells (*p*_*val*_ < 1*e* − 6, 1*e* − 9, Figure 6A) and lowly expressed in effector cells (*p*_*val*_ < 1*e* − 21, 1*e* − 9, Figure 6A), suggesting that they are required in the cycling state but not in the effector state, and that knocking them out could potentially block proliferation but not differentiation. We therefore re-analyzed proliferation data in perturbed T cells from the literature^36^ to check this hypothesis. From analyzing a genome-wide proliferation screen obtained by co-transducing OT-I CD8 T cells with *sgRegnase-1* and a genome-scale lentiviral sgRNA library,^36^ we found that the proliferation rate induced by perturbing our same target genes was positively correlated with the numbers of cells we observed in our data (ρ_*val*_ = 0. 45, *p*_*val*_ < 1*e* − 6, Figure 6B), suggesting that the prior proliferation data provides consistent evidence for the effects of our target genes. The two hits, *Ndufv2* and *Dimt1*, lie in the lower spectrum of all genes tested in the genome-wide screen^36^ in terms of promoting proliferation (bottom 88% and 84%) (Figure 6C). However, this trend was less pronounced when comparing with the population of target genes that overlapped with our two screens, which overall led to significantly less proliferation compared to all genes (*p*_*val*_ < 5*e* − 3, Figure 6C, Figure S7A).

**Figure 6.**
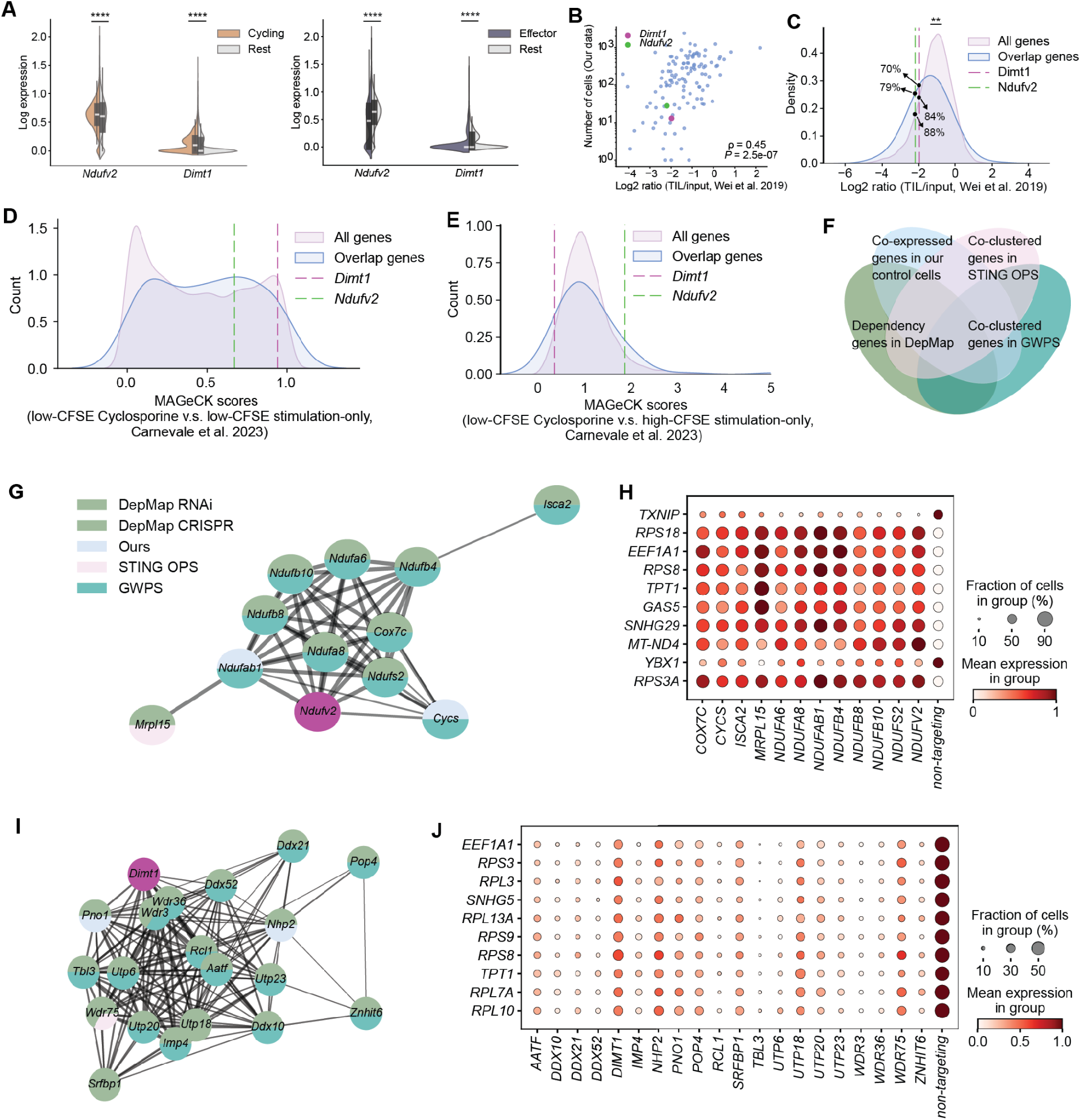
*Dimt1* and *Ndufv2* suppress proliferation but are resistant to blocking of activation. **(A)**, Violin plots overlaid with box plots indicating the expression levels of *Dimt1* and *Ndufv2* in cycling cells versus the rest of the cells (left) and in effector versus the rest of the cells (right) Differential expression measured by t-test with overestimated variance (*p*_*val*_ < 1*e* − 6, 1*e* − 9 for cycling cells, *p*_*val*_ < 1*e* − 21, 1*e* − 9 for effector cells). **(B)**, Scatterplot contrasting the numbers of cells observed for each knockout in our screens against the enrichments (log2(TIL/input ratio)) of the corresponding knockout in tumour-infiltrating OT-I cells relative to pre-transfer OT-I cells in a genome-scale CRISPR-Cas9 screen^36^ (Spearman ρ = 0. 45, *p*_*val*_ < 1*e* − 6). The top 2 hits from our second screen are indicated by colored dots. **(C)**, Distribution of enrichments (log2(TIL/input ratio)) in tumour-infiltrating OT-I cells relative to pre-transfer OT-I cells in a genome-scale CRISPR-Cas9 screen.^36^ All genes from this screen are shown in pink. Overlapping genes between this screen and our two screens are shown in blue. The comparison between all genes and overlapping genes are performed using one-side Mann-Whitney U test (*p*_*val*_ < 5*e* − 3).The top 2 hits from our second screen are indicated by dashed lines. **(D)**, Distribution of gene-level MAGeCK scores of dividing cells under immunosuppressive condition with Cyclosporine versus control in genome-wide CRISPR knock-out screens on human T cells.^55^ All genes from these screens are shown in pink. Overlapping genes with our two screens are shown in blue. The top 2 hits from our second screen are indicated by dashed lines. **(E)**, Similar as **(D)**, where the MAGeCK score is computed by comparing dividing cells treated with Cyclosporine against non-dividing cells in the control. **(F)**, Schematic of finding genes that are similar to *Dimt1* or *Ndufv2*. A gene will be considered if and only if it appears top in at least two lists out of the following criteria: co-expressed with the hit gene (*Ndufv2* or *Dimt1*) in our control cells, co-clustered with the hit gene in STING OPS^56^, co-dependent with the hit gene in DepMap RNAi or CRISPR screens^50^, or co-clustered with the hit gene in GWPS.^48^ **(G)**, STRING-Mus PPI^30,31^ between genes that are picked up by **(F)** with *Ndufv2*. The node color indicates which of the lists these genes were picked up from. **(H)**, Top 10 differentially expressed genes between cells that were perturbed by one of the connected genes in **(G)** and cells from non-targeting control in GWPS. **(I)**, Similar as **(G)** for *Dimt1*. **(J)**, Similar as **(H)** for *Dimt1*.

To assess the effect of perturbing these genes on cell activation, we leveraged data from genome-wide CRISPR knock-out screens in primary human T cells across multiple suppressive conditions,^40^ and found that both *Ndufv2* and *Dimt1* are significant hits in the screens (based on MaGeCK scores^55^) when comparing dividing cells (characterized by low carboxyfluorescein succinimidyl ester (CFSE) staining) with Cyclosporine versus dividing cells without cyclosporine (and T cell suppressing drug) (Figure 6D). Here, a higher MaGeCk score may suggest that loss of *Ndufv2* and *Dimt1* leads to enhanced cell division and/or resistance to the suppressive condition. However, the score may be high also if the denominator (i.e., dividing cells in the control setting without cyclosporine) is small. Therefore, we analyzed the MaGeCK score by comparing to non-dividing (characterized by high CFSE staining) stimulation-only: *Ndufv2* remained high, while *Dimt1* dropped to the lower spectrum in this score (Figure 6E). A similar behavior was observed when instead of Cyclosporine, Tacrolimus was applied [PMID 36002574] (Figure S7B-C). This indicates that the proliferation promoted by the depletion of *Dimt1*, a DepMap common essential gene,^50^ is more specific than *Ndufv2* depletion in suppressing proliferation in the context of agents such as Cyclosporine, Tacrolimus. However, it is worth noting that none of the MaGecK scores are correlated with proliferation directly (Figure S7D-G).

We next analyzed the pathways and downstream signatures of these two hits. For the pathway analysis, we obtained similar genes that are: (1) co-expressed with these two hits in our control cells, (2) co-clustered with these two hits based on perturbation-induced phenotypes in a genome-wide OPS screen of Stimulator of interferon genes (STING) trafficking,^56^ (3) dependency genes across the same cell lines as these two hits in DepMap,^50^ (4) co-clustered with these two hits in GWPS^48^ (Figure 6F, Methods, Supplementary Table 4). For genes that appeared in the top at least two times in the similarity search described above with respect to *Ndufv2* (Figure S7H-I), we retrieved a protein-protein interaction network from STRING^30,31^ (Figure 6G, Methods). We also analyzed the common differentially-expressed genes in response to perturbation of these genes in GWPS^48^ (Figure 6H, Methods). We found that *Ndufv2* is involved primarily in mitochondrial pathways which encode components of mitochondrial oxidative phosphorylation. It is possible that their disruption limits ATP production, ROS signaling, and mTOR–Myc activity, with some effector differentiation still initiated but selectively impaired while favoring the persistence of metabolically restrained progenitor-like T cells. However, their exact effect on anti-tumor immunity is not yet known. Similarly, we performed this analysis for *Dimt1* (Figure 6I-J, Figure S7J-K), and found that it controls ribosomal production and ribosome biogenesis and rRNA processing, and its disruption is expected to limit translational capacity, selectively impairing biosynthetically demanding effector differentiation while favoring the survival and accumulation of progenitor-like T cell states.

## Discussion

In this study, we analyzed the results of our Machine Learning Competition for Cancer Immunotherapy. Our inaugural competition was set up to test the state of the field in terms of algorithms for the prediction of gene-perturbation effects and was uniquely centered around a fundamental bottleneck in T cell engineering for cancer immunotherapy: the efficient identification of gene targets that modulate T cell fate.

In 2025, following our initial efforts, the Virtual Cell Challenge^25^ was launched with the similar goal to accelerate the development of algorithms to predict the effects of cellular perturbations. The Virtual Cell Challenge was particularly focused on gene expression, requiring participants to predict the effects of held-out single-gene perturbations in H1 human embryonic stem cells, and highlighted the difficulties in defining robust evaluation metrics based only on gene expression predictions.^57,58^ In contrast, we chose to center the prediction tasks around a well defined biological problem in a therapeutically relevant mouse model to accelerate the development of methods that not only succeed in predicting the transcriptional effect of held out perturbations, but can also predict the effect of unseen perturbations on cell state proportion changes. While most perturbation models prioritize predicting transcriptional responses, we found that predicting the biological objective directly, in our case T cell state proportions, outperformed models designed with an intermediate step of first predicting transcriptional responses. This suggests aligning model training closely with the ultimate downstream biological task can significantly improve prediction accuracy.

While the average transcriptional effect of gene perturbation at the single cell level may be subtle and the low-signal-to-noise ratio can easily overshadow the true biological signal of a perturbation, we believe that changes in cell state proportions are a more robust readout of the perturbation effect that is likely to be more impactful to drive new biological discoveries. In our competition, we tested the algorithms’ performance on two tasks, and found a clear difference in methods that performed well in predicting the effects of held out perturbations (Challenge 1) and those that did well in ranking the effect of unseen genes based on therapeutic objectives (Challenge 2). While the first task revealed which methods can predict well on average (based on the average prediction loss across all tested perturbations); the second measured primarily whether a method can distinguish which are the better targets amongst the top predicted ones, based on experimental validation of the top predictions. This second evaluation is potentially a more direct way to assess a method’s utility for nominating new targets with the potential to induce biologically relevant shifts in cell populations.

Our analyses showed that the the top-performing methods in both Challenges can be described using a similar two-step design, where in the first step, all genes are mapped to meaningful features constructed based on the provided data as well as existing online resources, and in the second step, these features are mapped to the predicted cell state proportion where the mapping is trained using the provided data. We found the first step to be a major separating factor between different methods’ performances, a result in line with recent studies^59,60^ that stress the importance of incorporating prior biological knowledge for perturbation effect prediction. Here, existing perturbational datasets constitute the strongest prior for predicting the effect of unseen perturbations, a finding confirmed by our own proposed gene features, which was curated from prior T-cell perturbation literature and performed best in our Challenge 1 benchmarks. However, the modest gain of the top methods in prediction accuracy (approximately 7.5% in L1 error) over the averaged perturbation baseline, which has been shown in literature^46^ to outperform many deep-learning models^61–65^, underscores the inherent difficulty of accurately predicting perturbation effects. Furthermore, the prediction error was highest for perturbations that resulted in minimal change from the non-targeting control, suggesting that current models struggle to accurately distinguish between a true null effect and a small-magnitude functional perturbation. The observed correlation between prediction error and the L1 distance from the training set’s mean effect emphasizes that the current models are primarily interpolating within the explored perturbation space. This indicates a fundamental need to expand the training set to encompass a broader spectrum of biological effects for a robust extrapolation to unseen genetic perturbations.

In our second Challenge, most of the algorithms-proposed genes had a lower objective score than the expert-selected 73 genes in our training screen. Nonetheless, the ML-guided approach picked up two targets *Dimt1, Ndufv2* with higher ICB and CAR-T objectives, which generally led to less terminal exhaustion and more progenitor states than all tested targets. Prior T-cell specific literature showed that their inhibition generally leads to less proliferation,^36^ while preventing the activation block induced by immunosuppressive drugs such as cyclosporine and tacrolimus.^40^ *Dimt1* is an essential gene controlling ribosomal functions, while *Ndufv2* encodes for a core subunit of mitochondrial Complex I that is involved primarily in mitochondrial respiratory chain and ROS production. Interestingly, the depletion of another gene involved in similar mitochondrial pathways as *Ndufv2, Ndufa4*, was recently found to be amplifying interferon signaling and promoting anti-tumor immunity in tumor-associated macrophages, as its repression increased mitochondrial DNA release into the cytoplasm and subsequent STING activation, thereby amplifying anti-tumor IFN-induced transcriptional programs in TAMs.^66^ The exact mechanism and impact of *Ndufv2* and *Dimt1* depletion on enhancing immunotherapy responses, remain an open question for future work.

Overall, our competition highlights both the promise and current limitations of machine learning for perturbation biology. Performance plateaus relative to simple baselines caution against overinterpreting predictive capacity in the absence of rigorous evaluation. At the same time, the identification of biologically meaningful targets demonstrates that structured benchmarking frameworks, with carefully defined objective functions, can yield experimentally actionable discoveries. Building on these insights, we established the Cell Perturbation Prediction Challenge (CPPC), an ongoing series of yearly competitions designed to continuously generate biomedically relevant perturbation datasets and provide an objective platform for iterative ML model testing and improvement. This iterative *experimentation-to-ML-to-experimentation* feedback loop is the necessary path forward to create robust, generalizable predictive models that can accelerate therapeutic design.

## Supporting information

Note S1. Implementation of methods and hyperparameter selection.

Table S1. List of the genes targeted in the CRISPR datasets.

Table S2. Top submissions in each of the 3 Challenges.

Table S3. Target genes ranked by objective functions.

Table S4. Crowd-sourced co-expressed / co-clustered / dependency genes with Ndufv2 and Dimt1.

## Supplemental Figures

**Figure S1.**
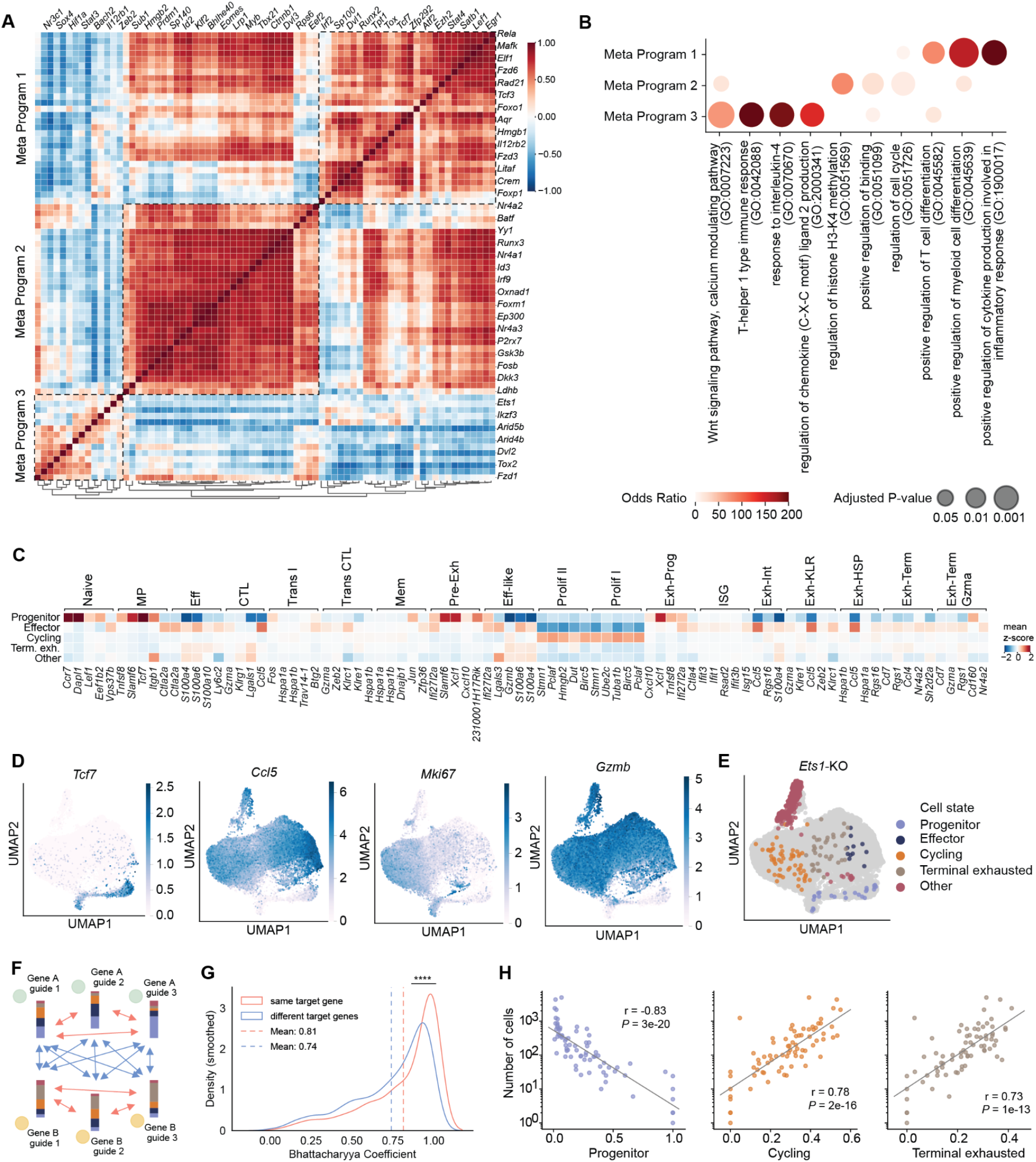
Further analysis of in-vivo CRISPR screen. **(A)**, Visualization of the correlation among the 73 knock-out genes based on the bulk gene expression profiles in perturbed cells. The gene target names are shown for the even columns on the top axis, and for the odd rows on the right axis. We identified three gene modules (Meta Programs) that induced similar transcriptional changes upon perturbation at the top2 levels of the dendrogram (bottom). **(B)**, Pathway annotations of the three meta programs. **(C)**, Heatmap showing the mean z-score for the top marker genes from previous literature’s annotated states^67^ in cells in each of our annotated states. **(D)**, UMAP visualization colored by marker gene expression. **(E)**, UMAP visualizations to highlight the cell state distribution for cells transduced with sgRNAs targeting *Ets1*, which is responsible for the emergence of the other cell states. **(F)**, Schematic showing how guides targeting the same/different genes are compared. For each pair of guides, a similarity coefficient is computed between the cell state proportions they induced. Then the coefficient between guides targeting the same gene is compared to coefficients targeting different genes. **(G)**, Histogram showing the distribution of the Bhattacharyya coefficients between same target gene versus different target genes, as illustrated in **(F)**. The coefficients between guides targeting the same gene are significantly higher than those between different guides (one-sided Mann-Whitney U test *p*_*val*_ < 1*e* − 4). **(H)**, Scatter plot showing the number of cells against the proportion of three states (Pearson *r* =− 0. 83, 0. 78, 0. 73, *p*_*val*_ < 1*e* − 19, 1*e* − 15, 1*e* − 13).

**Figure S2.**
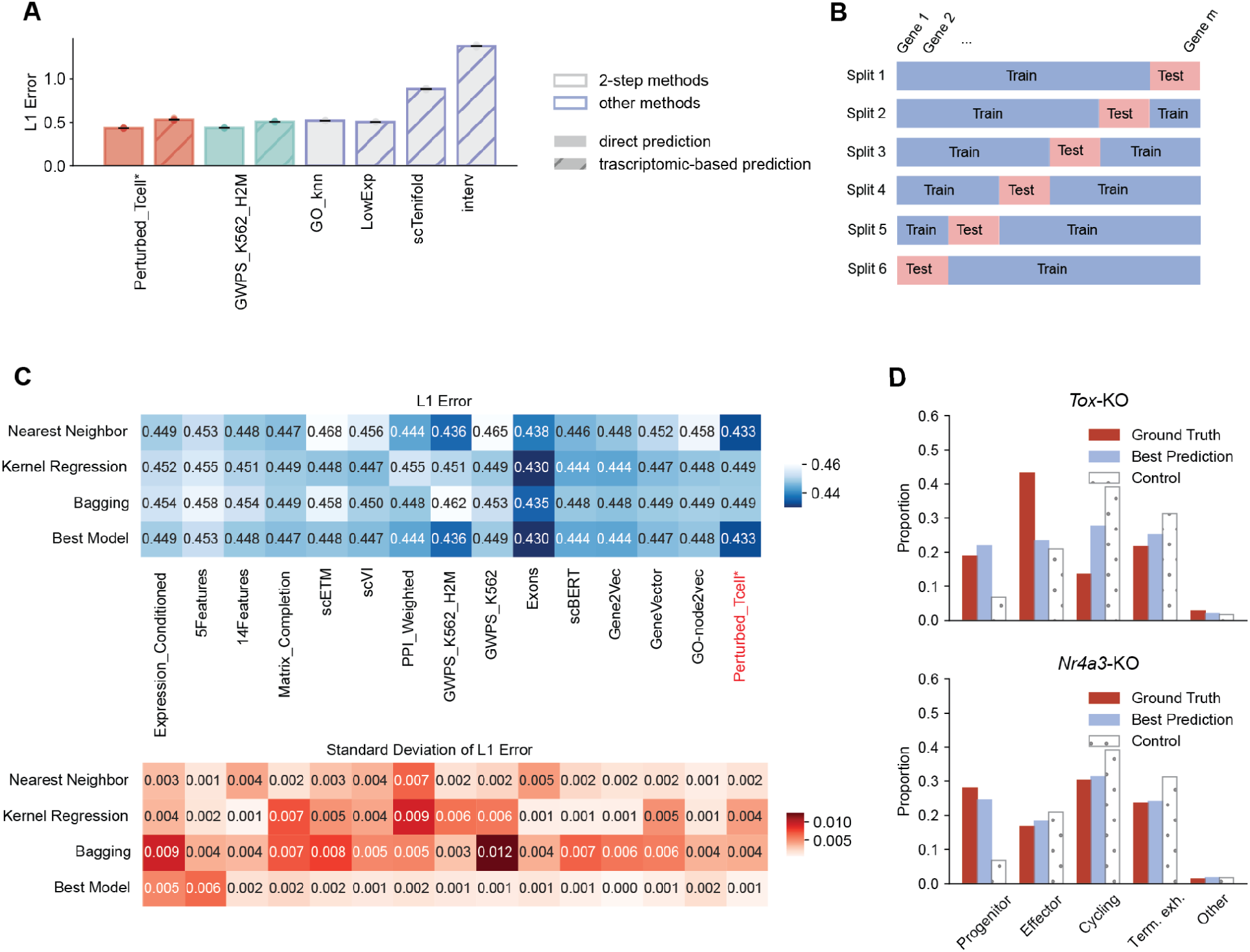
Further benchmarking results of methods to predict gene perturbational effect. **(A)**, *l*_1_ error comparing methods that can (grey boxes) versus cannot (blue boxes) be described by a 2-step procedure. The shading indicates if a method directly predicts cell state proportion vectors (direct, unshaded bars) or if a method first predicts the transcriptomic distributions and then classifies each cell into cell states (2step, shaded bars). **(B)**, Schematic showing the train-test splits and evaluation framework of the benchmark. **(C)**, *l*_1_ errors of different methods, across combinations of gene features and predictors (first three rows). The last row shows the best predictor of each gene feature from the first three rows. Top plot shows the averaged *l*_1_ error, where the bottom plot shows the standard deviation. **(D)**, Barplots of the prediction from the best method, the control distribution, and ground-truth proportion of two example genes.

**Figure S3.**
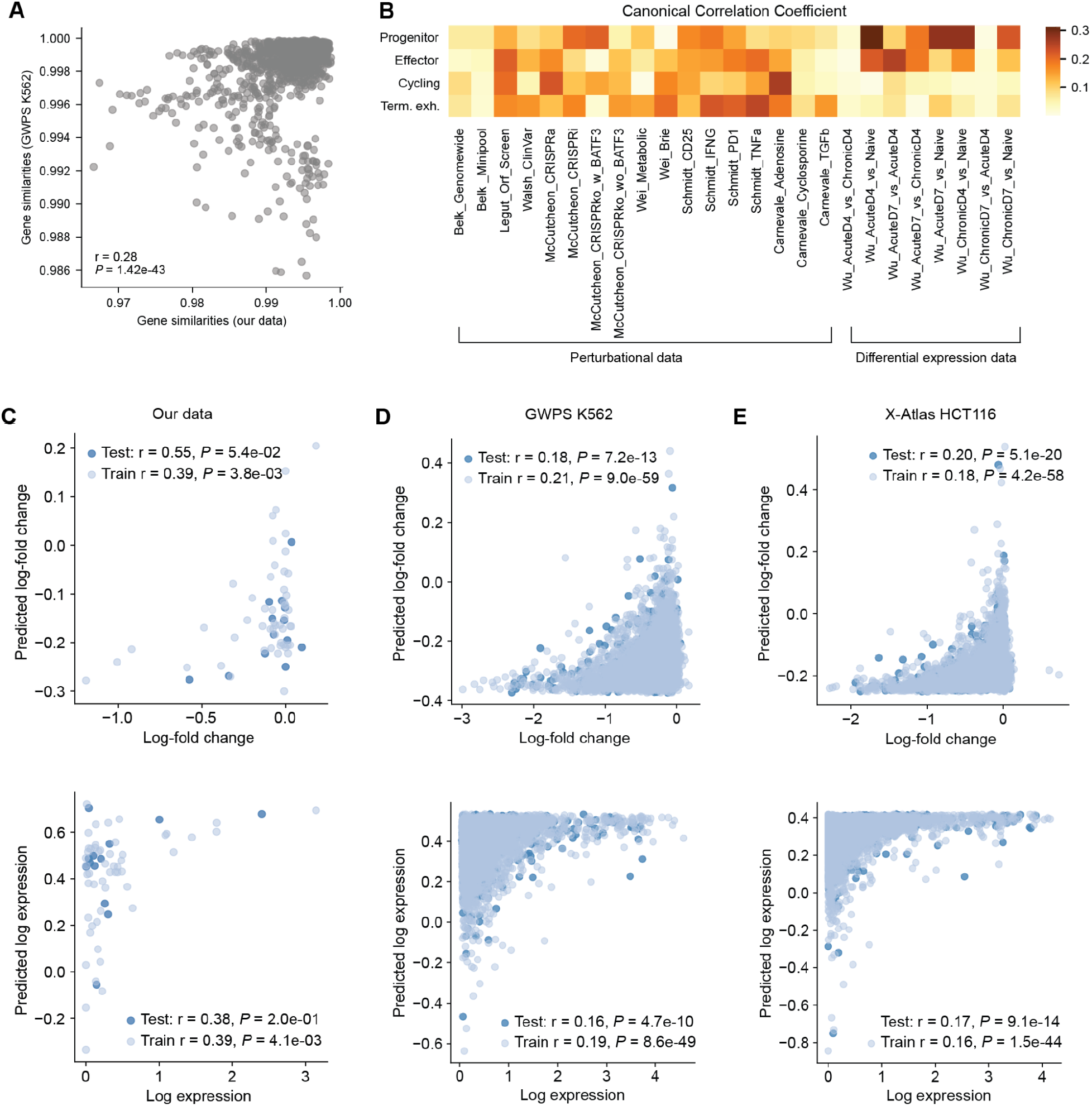
Feature analysis of top 3 methods in the benchmark. **(A)**, Scatterplot comparing the cosine similarities of perturbational changes of pseudo-bulk expression between pairs of genes in our dataset and that in GWPS.^48^ **(B)**, Canonical correlation between the proportions of 4 cell states and different gene features from the datasets that we used in the Perturb_Tcell method. **(C-E)**, Evaluation of using exon features to predict target gene log-fold changes between control and perturb (top rows) and log gene expression in the control (bottom rows) in our data **(C)**, in GWPS^48^ **(D)**, and in X-atlas on HCT116 cell line^51^ **(E)**.

**Figure S4.**
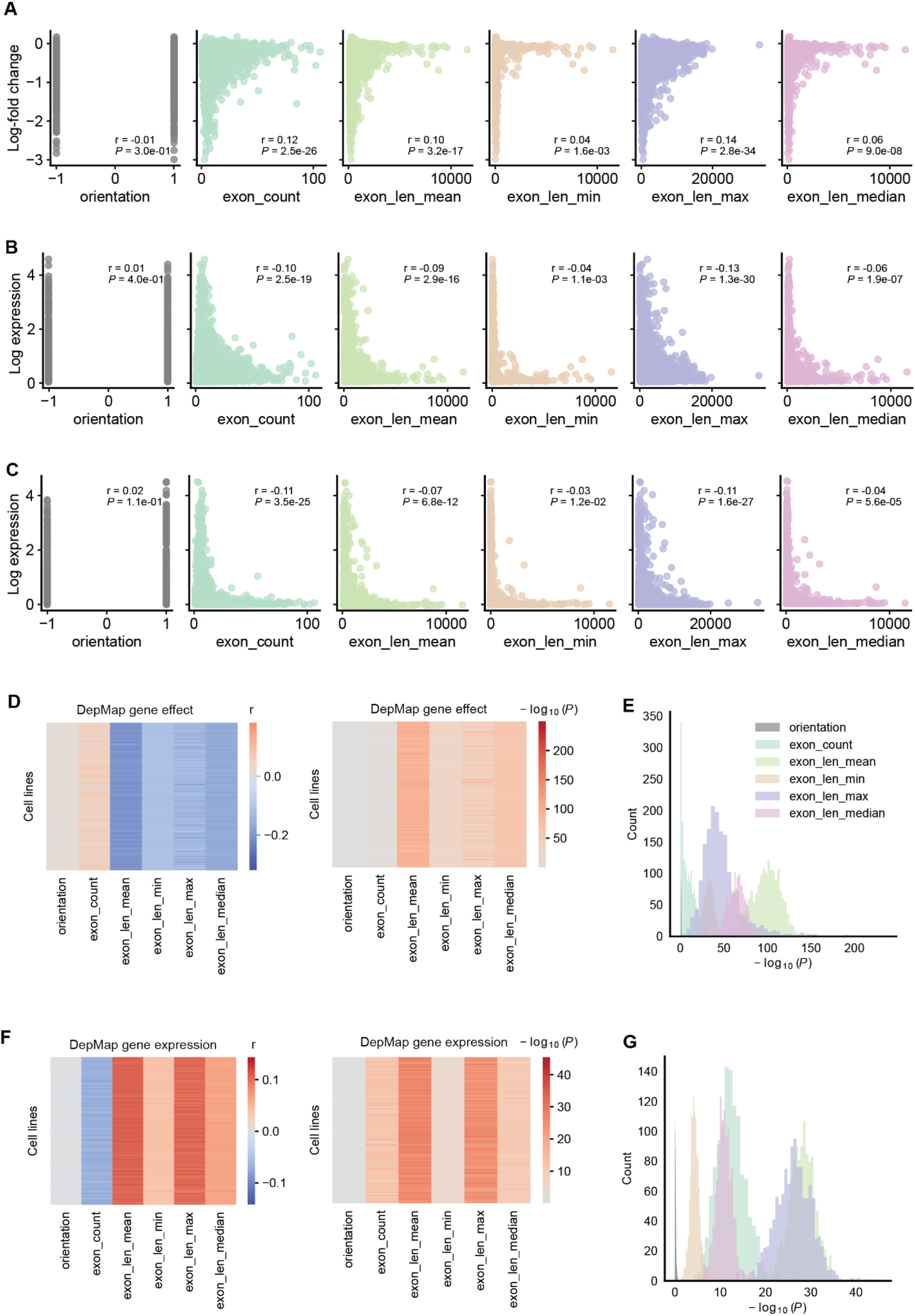
Analysis of individual exon features. **(A-C)**, Correlation between different exon features and log-fold changes of the gene comparing control and perturbation targeting the corresponding gene **(A)**, or log-expression of the gene in the control distribution in GWPS^48^ **(B)**, or in a single-cell RNA-seq of PBMC with no perturbations^49^ **(C)**. **(D-G)**, Correlation between different exon features and the effects of knocking out the corresponding genes cross cell lines **(D-E)**, or bulk log expression of the corresponding genes across cell lines in DepMap^50^ **(F-G)**. Heatmaps show the correlation (left) and p-values (middle). Histograms show the distribution of p-values (right).

**Figure S5.**
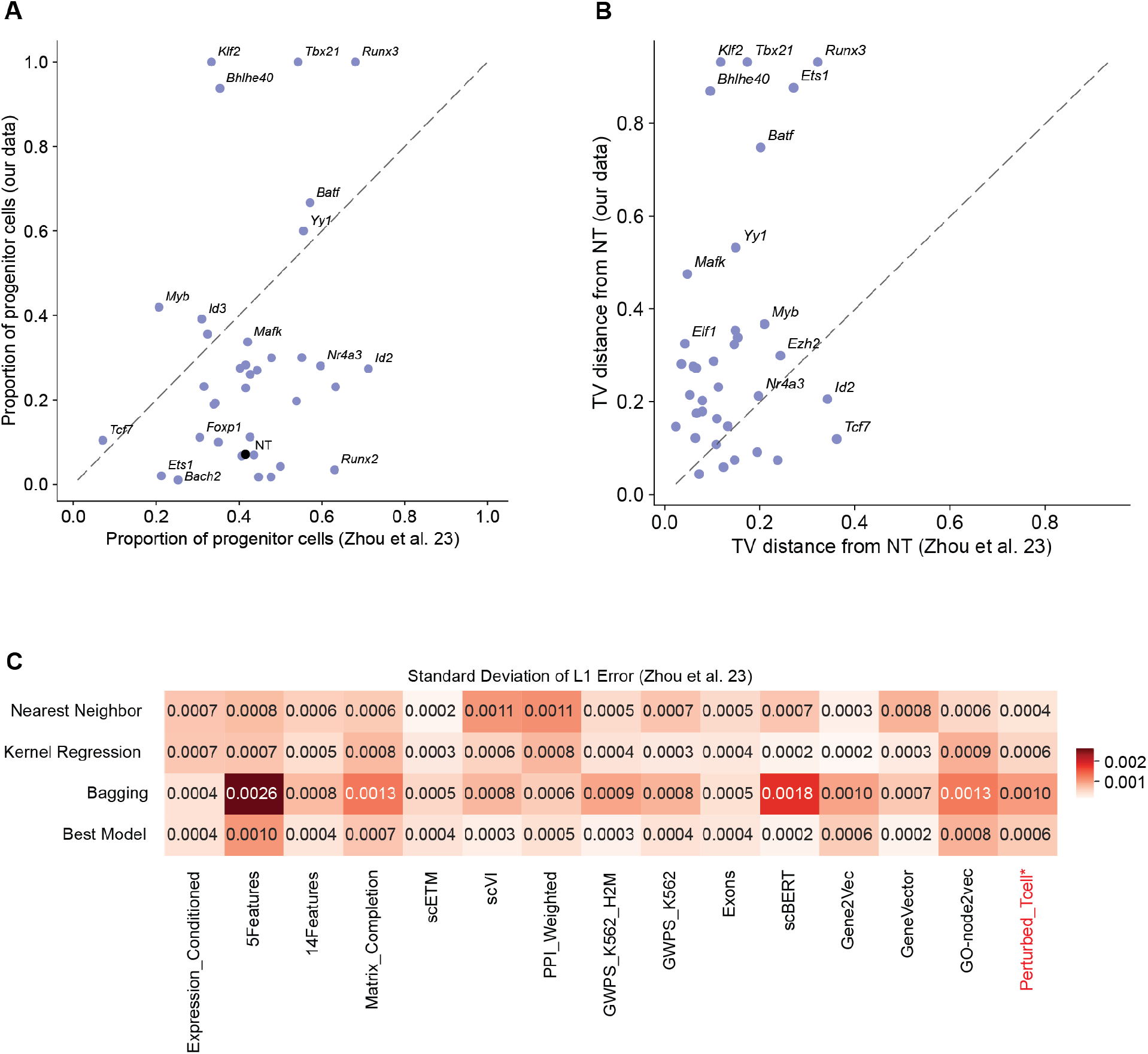
Further analysis of the public in-vivo Perturb-seq. **(A)**, Scatterplot comparing the progenitor proportion of genes that were both perturbed in our dataset and the public Perturb-seq data^53^ (one-side Mann-Whitney U test with *p*_*val*_ < 1*e* − 4). **(B)**, Scatterplot comparing the total variation distances between the perturbation induced cell state proportions to the control cell state proportions on genes that were both perturbed in our dataset and data from the public Perturb-seq data^53^ (one-side Mann-Whitney U test with *p*_*val*_ < 1*e* − 4). **(C)**, Standard deviation of L1 errors (whose averages are shown in Figure 3E) of different methods, across combinations of gene features and predictors (first three rows). The last row shows the best predictor of each gene feature from the first three rows.

**Figure S6.**
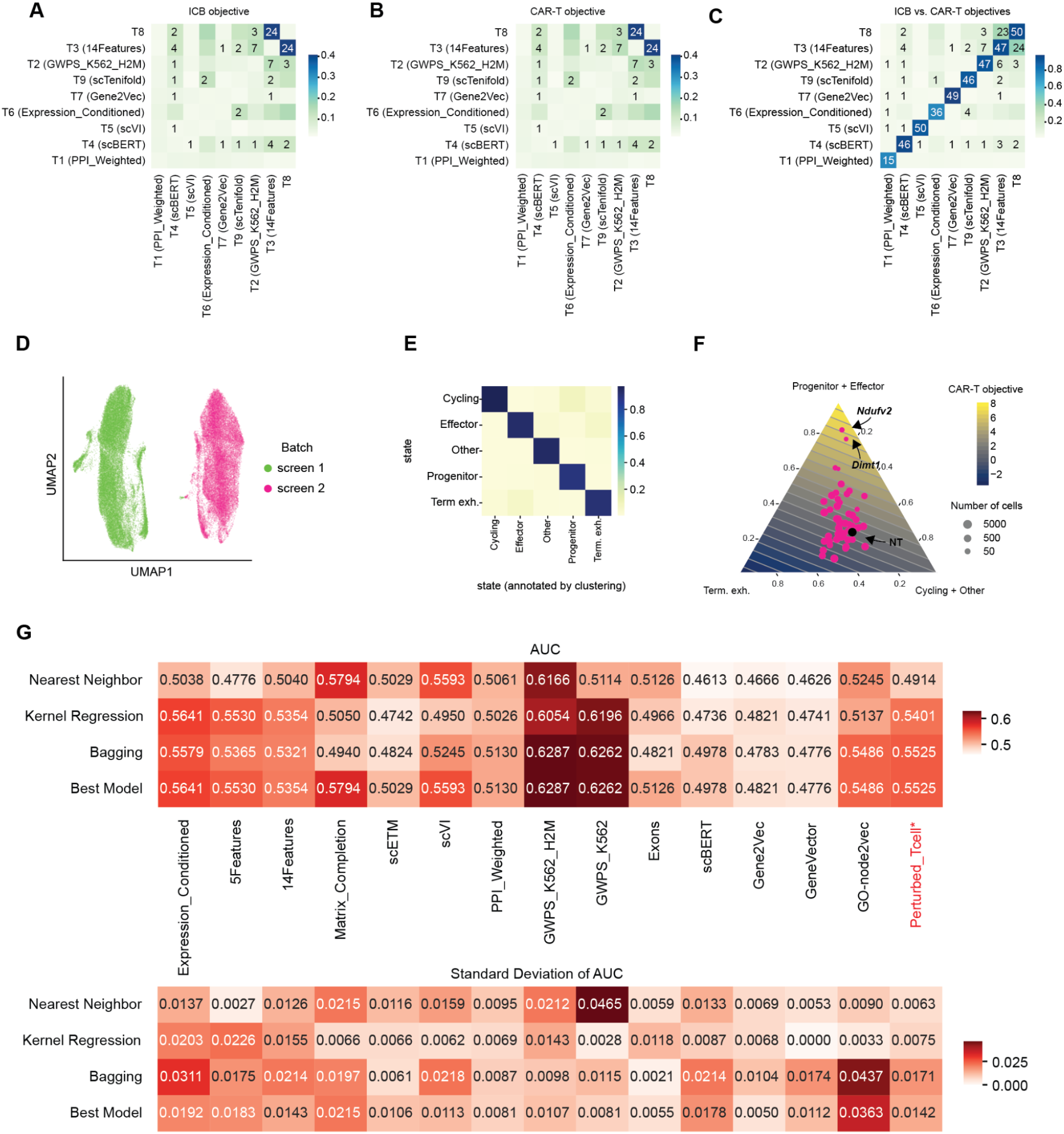
Further analysis of newly proposed genes and the validation screen. **(A-B)**, Overlap between top 50 genes ranked by each team for the ICB objective **(A)** and CAR-T objective **(B)**. Coloring indicates the area under the curve between two rank lists and annotation indicates the number of overlapping genes. Diagonals are manually set to zero, as it corresponds to comparing the same list against itself. **(C)**, Overlap between top 50 genes ranked by each team in terms of ICB objective versus in terms of CAR-T objective. Color and annotation are the same as **(A-B)**. **(D)**, UMAP visualization of the validation screen (screen 2) and the first screen (screen 1) before batch correction. **(E)**, Confusion matrix of cell state annotation versus re-annotation based on the clustering-comparison approach on the first screen. **(F)**, Ternary plot visualizing perturbations in our second screen in terms of the CAR-T objective. The heatmap coloring indicates the smoothed averaged objective function values of the proportions at the corresponding spot. Each dot corresponds to a perturbation (or NT, as annotated by the black dot), where the size of the dots indicates the number of cells for that perturbation and top 2 perturbations are annotated. **(G)**, Validation of methods when trained only on screen 1 to rank the perturbations screen 2 according to the CAR-T objective. Top plot shows the area under the curve when comparing the predicted order to ground-truth order, averaged across 5 runs. Bottom plots show the standard deviation across 5 runs.

**Figure S7.**
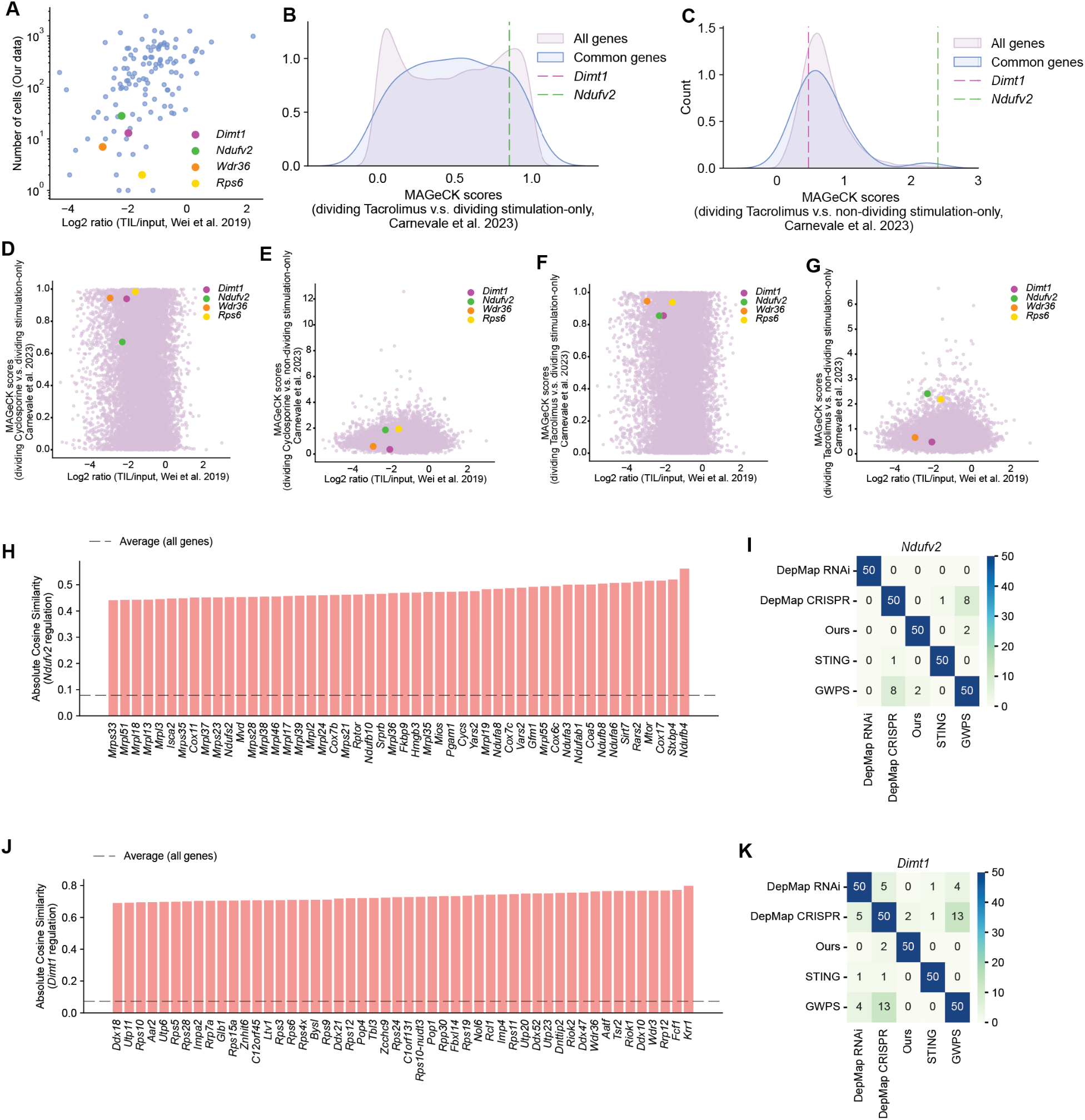
Further analysis of *Dimt1* and *Ndufv2*. **(A)**, Same as Figure 6B but with additional annotation on *Wdr36* and *Rps6*, which are genes that potentially lie in similar pathways as *Dimt1*, as identified in Figure 6F. **(B-C)**, Similar to Figure 6D-E, but on cells treated with Tacrolimus. **(D-G)**, Scatterplot contrasting a gene in terms of enrichment from Figure 6c and division under immunosuppressive conditions from Figure 6D-E and **(B-C)**. **(H)**, Bar plot showing the top 50 genes that co-expressed with *Ndufv2* in GWPS.^48^ The measure is the absolute cosine similarity (a measure of co-expression) to *Ndufv2* of the changed pseudobulk expression comparing perturbed cells and control cells. The dashed line indicates the average absolute cosine similarity to *Ndufv2* across all genes. **(I)**, Heatmap showing overlap between top 50 genes in lists of the criteria described in Figure 6F. **(J-K)**, Similar as **(H-I)**, for *Dimt1*.

## RESOURCE AVAILABILITY

### Lead contact

Further information and requests for resources and reagents should be directed to and will be fulfilled by the lead contact.

### Data and code availability

Data: The datasets produced by this paper can be accessed at GSE327731. Code: All original code is publicly available and has been deposited at: https://github.com/uhlerlab/cancer_immunotherapy_data_science_challenge

## ACKNOWLEDGEMENTS

We would like to thank all the participants in the Challenge, partially listed below under CPPC Community, for their algorithms and active involvement in the project. We acknowledge E. Forte for valuable discussions and editing the manuscript. Primary financial support for the Challenge came from the Eric and Wendy Schmidt Center. J.Z. was partially supported by the Eric and Wendy Schmidt Center at the Broad Institute and by an Apple AI/ML PhD Fellowship. Y.P. was supported by NIH/NIAID grant DP2AI171161, NSF CAREER grant 2238831, Ludwig Institute for Cancer Research, Weill Cancer Hub East. O.A. was supported by funds from the Klarman Cell Observatory and the Eric and Wendy Schmidt Center at the Broad Institute. N.H. was supported by the Mark Foundation Endeavor Award. C.U. was partially supported by NCCIH/NIH (1DP2AT012345), NIDDK/NIH (5RC2DK135492-02), ONR (N00014-24-1-2687), and the United States Department of Energy (DE-SC0023187).

## CPPC Community

we established the Cell Perturbation Prediction Challenge (CPPC), an ongoing series of yearly competitions designed to continuously generate biomedically relevant perturbation datasets and provide an objective platform for iterative ML model testing and improvement. Below are the participants, grouped by teams (affiliation are collected by the time that challenges concluded), who won prizes for the cancer immunotherapy challenges described in this work:

- Marios Gavrielatos and Konstantinos Kyriakidis, Greece
- Yuzhou Gu, Anzo Teh, Yanjun Han, and Brandon Wang, U.S. (MIT)
- Peter Novotný, Poland
- Brody Langille, Jordan Trajkovski, and Elizabeth Hudson
- Marc Glettig
- Ai Vu Hong, researcher at Genethon, France
- Saket Kunwar, independent researcher, Nepal
- John Gardner, freelance data scientist
- Basak Eraslan, postdoctoral researcher holding a joint position at the Regev Lab in Genentech and Kundaje Lab at Stanford University
- Haoyue Dai, Kun Zhang, Ignavier Ng, Yujia Zheng, Xinshuai Dong, and Yewen Fan from Carnegie Mellon University; Petar Stojanov, postdoctoral fellow at the Eric and Wendy Schmidt Center; Gongxu Luo, Mohamed bin Zayed University of Artificial Intelligence; and Biwei Huang, University of California, San Diego
- Liu Xindi, freelance programmer
- Johnson Zhou, Camille Sayoc, and Yi-Cheng Peng, Master’s students of the Faculty of Engineering and IT at the University of Melbourne, Victoria, Australia
- Dariusz Brzeziński and Wojciech Kotlowski from Poznań University of Technology in Poland
- Salil Bhate, postdoctoral fellow at the Eric and Wendy Schmidt Center
- Irene Bonafonte Pardàs, Artur Szalata, and Benjamin Schubert from Helmholtz Center Munich and Miriam Lyzotte from Mila - Quebec AI Institute

## AUTHOR CONTRIBUTIONS

J.Z., M.S., O.A., N.H., and C.U. conceived and designed the research. Y.P. contributed to the early design of the ML challenge. J.Z. performed the data analysis and implemented the algorithms. M.S., M.M., and O.O. performed the experiments. O.A., N.H., and C.U. supervised the study. All authors wrote the paper.

## Methods

### In vivo pooled CRISPR screens with scRNA-seq

#### sgRNA library design and cloning into pRDA_206_Thy1.1

The sgRNA oligonucleotides were designed using the Broad CRISPR algorithm: https://portals.broadinstitute.org/gppx/crispick/public. Pooled CRISPR sgRNA libraries were constructed by cloning pooled sgRNA containing oligonucleotides via golden gate assembly into pRDA_206_Thy1.1, a lentiviral sgRNA expression vector containing a U6 promoter upstream of the sgRNA cassette and a PGK–Thy1.1 surface marker to enable enrichment of transduced cells. Cloned plasmid DNA was delivered to Endura electrocompetent cells via electroporation, and plasmid DNA was extracted after overnight culture using Zymopure plasmid maxiprep kits.

#### Lentiviral production

Lentivirus was produced by transient transfection using Lipofectamine 3000 of HEK293T cells with the pooled pRDA_206_Thy1.1 sgRNA library plasmid together with psPAX2, pVSVG, and pCAG-ECO. Viral supernatant was harvested 48 hours after transfected, spun at 200×G and filtered, then aliquoted and stored at −80 °C until use. Functional titer was assessed directly on primary T cells to select an input that achieved the desired transduction rate.

#### Naïve CD8 T cell isolation and lentiviral transduction

Naïve CD8 T cells were isolated from spleens of H11-Cas9 × OT-I mice (6–16 weeks old, male and female) using magnetic bead–based enrichment for naïve CD8 T cells. Purified naïve CD8 T cells were transduced with pooled lentivirus on retronectin-coated plates in complete T cell medium supplemented with IL-7 and IL-15 at 10 ng/mL. Viral input was titrated on T cells to achieve 20–40% Thy1.1 positivity, consistent with predominantly single-perturbation transduction in pooled screens. Cells were maintained in IL-7/IL-15–supplemented medium for 5 days post-transduction without TCR activation.

#### Enrichment of transduced cells and adoptive transfer

On day 5 post-transduction, Thy1.1+ cells were enriched by magnetic selection to generate a transduced donor population. Enriched Thy1.1+ OT-I Cas9+ CD8+ T cells were adoptively transferred into congenic recipient mice by tail vein injection (5 × 10^5^ cells per mouse). Congenic markers varied by experiment; donor and recipient combinations (CD45.1/CD45.2) together with Thy1.1 expression were used to enable identification of transferred T cells by flow cytometry. T cells were transferred one day prior to tumor implantation.

#### Tumor implantation and monitoring

Recipient mice were implanted subcutaneously with B16-OVA melanoma cells (5 × 10^5^ cells per injection) in 200 µL PBS into bilateral flanks. Tumor growth and general health were monitored in accordance with animal welfare guidelines.

#### Tumor harvest and isolation of tumor-infiltrating CD8 T cells

Tumors were excised, minced, and dissociated using Miltenyi tumor dissociation kits to recover tumor-infiltrating lymphocytes. Dissociated material was filtered to remove debris, and lymphocytes were enriched by density gradient separation with LympholyteM (Cedarlane). CD8 T cells were subsequently purified from tumor infiltrates using Miltenyi magnetic bead–based CD8 enrichment. Only tumor-derived T cells were used for downstream single-cell profiling.

#### Hashtag staining, single-cell library preparation and sequencing

In the following, the training Perturb-seq screen (Figure 1) and the validation screen (Figure 4) are referred to as Screens 1 and 2, respectively. To enable multiplexing of samples from individual mice in Screen 2, purified tumor-infiltrating T cells from each mouse were stained with BioLegend mouse TotalSeq-C hashtag antibodies according to the manufacturer’s protocol, including stringent washing steps to minimize ambient oligonucleotide background. Hashtag-labeled samples were pooled and processed using the 10x Genomics Chromium 5′ single-cell gene expression workflow with Feature Barcode chemistry for cell-surface antibody–derived tags and direct CRISPR guide capture. Libraries were prepared following the manufacturer’s instructions to generate: (i) 5′ gene expression libraries, (ii) feature barcode libraries for hashtags and cell-surface proteins when appropriate, and (iii) guide capture libraries reporting sgRNA identity. Final libraries were sequenced on an Illumina NextSeq platform using 10x Genomics–recommended read configurations and target sequencing depths for 5′ gene expression, feature barcode, and direct guide capture libraries.

### Data preprocessing and normalization

Screens 1 and 2 were processed and normalized separately using the same pipeline. Readouts from the *in vivo* CRISPR screens were processed with Cell Ranger^68^ to generate raw count matrices. After merging all sequencing lanes within each screen, we obtained 71,393 cells for Screen 1 and 75,227 cells for Screen 2. Quality control was performed to remove low-quality cells that either (i) received no perturbation or multiple perturbations, or (ii) had elevated mitochondrial read fractions indicative of stressed or dying cells.

To reduce technical noise, relatively stringent thresholds were applied. For Screen 1, cells were removed if they met any of the following criteria: fewer than 5 sgRNA UMIs; a ratio of the second-most abundant sgRNA to the most abundant sgRNA greater than 40%; a ratio of the most abundant sgRNA less than 60%; or a number of detected features (as called by Cell Ranger^68^) not equal to one. Cells with more than 5% mitochondrial counts were also removed. After filtering, 31,009 cells remained.

For Screen 2, additional antibody hashtagging was performed on individual mice prior to pooling. The same thresholds as in Screen 1 were applied for perturbation assignment, and cells with zero or more than one antibody hashtag were additionally removed. Cells with more than 10% mitochondrial counts were initially filtered; however, this threshold resulted in a distinct Leiden cluster with high mitochondrial content, which was subsequently removed. After quality control, 25,738 cells remained.

For each screen, the filtered raw count matrices were normalized by scaling the total counts per cell to 5,000, followed by a *log*(*x* + 1) transformation. When considering perturbations for the benchmarks, perturbations represented by very few cells were excluded due to large variance in annotated cell state proportions. A minimum cell number cutoff was therefore applied: 32 cells for Screen 1 and 10 cells for Screen 2, reflecting the different numbers of cells passing quality control in each screen.

We also analyzed a public screen^53^ when benchmarking the methods. For this analysis, we directly used the dataset after quality control, including *Tox*− cells and a perturbation-specific cluster that were excluded for their intratumoral CTL developmental trajectory (see Methods in their paper^53^). We then normalized the raw counts of this dataset using the same procedure described above.

### Cell state annotation

Normalized data from the previous section were used to annotate cell states. For Screen 1, *de novo* clustering was performed, and clusters were annotated using canonical markers. For Screen 2 and the public screen, a supervised approach was applied, leveraging annotations from Screen 1.

For Screen 1, dimensionality reduction to the top 50 principal components (PCs) was performed, followed by construction of a nearest-neighbor graph using 30 neighbors. Leiden clustering was applied with a resolution of 0.4, and each cluster was annotated by comparing its differentially expressed genes to known T cell state markers from the literature (Figure S1C-D). Hyperparameters were chosen based on qualitative assessment of the strength and separation of cluster-specific signatures.

For Screen 2, batch correction with Screen 1 was performed by (1) subsetting to shared genes, concatenating the normalized count matrices, (2) performing dimensionality reduction to the top 50 PCs, and (3) applying Harmony with default parameters to obtain batch-corrected PCs.^54^ Using these PCs, a nearest-neighbor graph was constructed with 30 neighbors, Leiden clustering was applied with a resolution of 1.5, and each cluster was assigned to a cell state based on the majority vote of cells previously annotated in Screen 1.

For the public screen^53^, we used the same procedure as for Screen 2 to annotate cell states, with the exception that we applied Harmony twice since there were still batch effects after the first application and that certain clusters were not annotated with the five states. This is because a few cells in their dataset were not observed in our dataset (Figure 3B), which is possibly a result of the experimental differences described in the main text. We excluded these cells when computing the cell state proportions for benchmarking the methods.

### Challenge setup and results

We collaborated with Topcoder to host the challenge. More than 1,000 participants from 87 countries, with backgrounds in mathematics, computer science, biology, and related fields, registered for the competition. To provide the necessary background, we developed introductory lectures on cancer immunotherapy and perturbation-based single-cell genomics^69^. The challenge tasks were designed to support experimental design, as explained in the main text. Because Challenge 2 relies on the objective functions defined in Challenge 3, we provide the details of the challenges below in the order: Challenge 1, 3, and 2.

In Challenge 1, seven randomly selected knockouts targeting *Aqr, Bach2, Bhlhe40, Ets1, Fosb, Mafk*, and *Stat3* were held out. Participants were asked to predict the corresponding five-dimensional vectors of cell state proportions. Denote the ground-truth and the predicted cell state proportions for perturbation *i* as 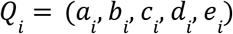 and 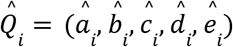 respectively.

The absolute error is defined as

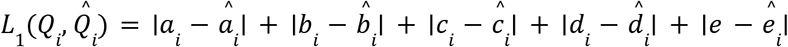

Three held-out knockouts targeting *Aqr, Bach2*, and *Bhlhe40* were used as the validation set, for which participants were allowed three submissions and received feedback on their averaged absolute error over the validation set. The final ranking was based on the average absolute error over the remaining four held-out knockouts, with one final submission permitted. The winners of Challenge 1 and their submitted methods are listed in Table S2. Predictions from these winners were used to select targets for Challenge 2.

In Challenge 3, participants were asked to propose scoring functions that rank perturbations according to their ability to shift cells toward a desired state. More precisely, the scoring function should take as input the gene expression distribution *P*_*i*_ resulting from knockout of gene *i*, the gene expression distribution *P*0 of unperturbed cells, a target cell state proportion vector *Q*, and output a scalar score for perturbing *i*. Higher scores indicate perturbations whose resulting cell state distributions are closer to *Q*.

Two representative objectives were provided: the ICB objective and the CAR-T objective, motivated by desired cell states in ICB and CAR-T therapies, as described in the main text. Both objectives operate on summary statistics rather than directly on *P*_*i*_ and *P*_0_ . Specifically, let 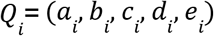 and 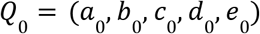 denote the cell state proportion vectors for knockout *i* and unperturbed cells, respectively, where proportions correspond to (progenitor, effector, terminally exhausted, cycling, other) cell states. The ICB objective was defined as

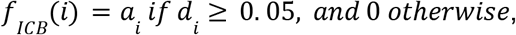

and the CAR-T objective as

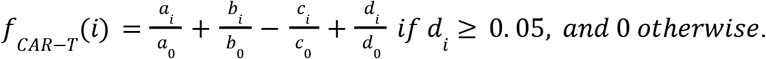

Note that the unperturbed cell state proportion vector in the training Perturb-seq screen (Screen 1) was *Q*_0_ = (0. 0675, 0. 2097, 0. 3134, 0. 3921, 0. 0173). Therefore the optimal cell state proportion, i.e., the target cell state proportion *Q*, for both objectives is *Q* = (0. 95, 0, 0, 0. 05, 0), where the two objectives measure the gap between *Q* _*i*_ and *Q*_0_ differently. The ICB objective prioritizes progenitor cell enrichment, whereas the CAR-T objective upweights effector and cycling states while penalizing terminal exhaustion.

Based on these two representative objectives, participants were asked to propose alternative objectives in a written report. The submissions were evaluated via a single-blind peer review followed by meta-review. Reviewers were recruited from volunteer participants; each reviewer evaluated a batch of three randomly selected submissions excluding their own, provided a brief summary, and ranked their three assigned submissions. The final ranking across all submissions was obtained by averaging reviewer ranks, followed by our review of the top submissions. The top three winning submissions with author consent are reported in Table S2. The winning objective defined the score for perturbation _*i*_ as

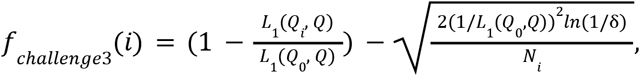

where *N*_*i*_ is the number of cells observed with knockout *i* and δ is a hyperparameter chosen based on the allowed estimation error (typically 0. 05). The first term equals 1 when *Q* _*i*_ = *Q*, is non-negative when the knockout performs at least as well as the unperturbed condition, and is negative otherwise. The second term provides a lower confidence bound correcting for finite-sample uncertainty, derived from the Hoeffding inequality, with range parameter 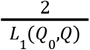.

In Challenge 2, participants were asked to predict the resulting five-dimensional vector of cell state proportions for all 15,077 expressed genes measured in Screen 1 that were not among the targets already tested. Based on the predicted proportions, the 15,006 previously untested genes (15,006 = 15,077-71 as 71 of the 73 genes were measured in expression) were ranked using two objective functions, *f*_*ICB*_ and *f*_*CAR-T*_, defined above, with the knockout cell state proportion *Q* _*i*_ replaced by the predicted proportion 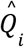. This resulted in two ranked gene lists. We did not rank submissions’ predictions using the top-performing objective *f*_*challenge3*_ from Challenge 3, as explained in the main text, because that objective requires the additional input *N* _*i*_, which was not included in the participants’ predictions.

We experimentally tested the top-ranked genes from the top submissions of Challenge 1 for both objectives,*f*_*ICB*_ and *f*_*CAR*−*T*_, in the validation screen (Screen 2). The experimentally validated genes were then ranked using the same objectives; for the CAR-T objective, we adapted the scoring by computing *Q*_0_ from the validation screen when evaluating *f*_*CAR-T*_. For each objective, we compared the ground-truth ranking of validated genes to the predicted ranking using an area-under-the-curve (AUC) metric, defined as follows. Let the genes ranked in the validation screen be [*g*_1_, …, *g*_*K*_], and let the predicted ranking be [*g* _*σ* (1)_, …, *g* _*σ* (*K*)_]. For *k* = 1, 2, …, *K*, we plotted the fraction of genes in [*g*_1_, …, *g*_*k*_] that appeared in [*g* _*σ* (1)_, …, *g* _*σ* (*k*)_], i.e.,

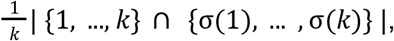

on the y-axis against the normalized list size *k*/*K* on the x-axis, and computed the area under the curve over the interval [0, 1]. This AUC score lies between 0 and 1 and has an expected value of 0. 5 for a random ranking, corresponding to points along the diagonal.

We also included *f*_*challenge3*_ in this comparison. For this objective, the ground-truth ranking was computed using the observed number of cells for each perturbation, whereas the predicted ranking used a version of the objective without the uncertainty term that depends on cell counts. For each submission, we computed AUC scores for all three objectives.

This evaluation for objectives *f*_*ICB*_ and *f*_*CAR-T*_was performed on both the filtered Screen2, in which perturbations with low cell numbers were removed as described in data preprocessing and normalization above, and the unfiltered Screen2, in which perturbations failing the minimum cell number cutoff were assigned a score of zero. For objective *f*_*challenge3*_, we evaluated it using just the unfiltered Screen2, as the additional uncertainty term captures the information about cell numbers. We then averaged the three scores for *f*_*challenge3*_, and *f*_*ICB*_, *f*_*CAR-T*_on filtered Screen 2, and separately averaged the three scores for *f*_*challenge3*_, and *f*_*ICB*_, *f*_*CAR-T*_on unfiltered Screen 2. The final score for each submission was defined as the minimum of these two averaged scores to ensure robustness. The winning submissions and their submitted methods are listed in Table S2.

### Implementation of top methods and our own proposed feature

We selected the top 20 methods from Challenges 1 and 2 for further analysis. To enable benchmarking across different train-test splits, we reimplemented these methods based on the submitted reports and code (Table S2). Because most of these methods can be summarized as a two-step mapping, we reimplemented them using the following formulation.

The prediction task can be formulated as learning a mapping *m* that takes as input a target gene *i* and outputs a 5-dimensional vector *Q*_*i*_ representing the proportions of the five cell states amongst the cells where gene *i* is knocked out, in other words, *m*(_*i*_) = *Q* _*i*_ . Screen 1 provides this mapping for its target genes, denoted as set *E*_1_ . To predict the effect of a perturbation not tested experimentally, a computational model must be learned to emulate *m* using Screen 1, where *m* can output meaningful *Q*_*i*_ for *i* not in *E*_1_ . A general way to obtain such *m* can be considered as a two-step mapping: (1) map each possible target gene *i* into a feature representation *r*(_*i*_) = *R*_*i*_, using either the provided experimental data or existing databases or both, (2) then fit a model *g* mapping *R*_*i*_ to *Q*_*i*_ on Screen 1. In other words, we consider the mapping of features *m* of form

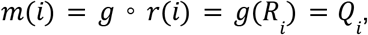

where *r* should be able to map all possible perturbations *i* to meaningful *R*_*i*_ using any existing datasets and information, and the proportion of gene features *g* is learned using pairs of (*R*_*i*_, *Q*_*i*_) for *i* in *E*_1_, where *Q*_*i*_ is measured (Figure 2A).

This two-step mapping allows us to combine gene features proposed by different teams with predictors proposed by other teams, and to disentangle the contributions of the gene feature choice in the first step from the predictor used in the second step. For the first step of constructing *r*, we derived 14 distinct gene features from the top submissions (Figure 2B), which are described in detail in Note S1. There we also described our own curated features, generated by integrating prior perturbed data on T cells, resulting in a total of 15 gene features For the second step of constructing *g*, a variety of predictors were employed by the top methods. We distinguished predictors by whether they directly predict cell state proportions or first predict transcriptomic profiles and subsequently infer proportions. For the direct approaches, we implemented three representative predictors: a simple and interpretable baseline (nearest neighbor), a canonical nonlinear method (kernel regression), and an ensemble method (bagging). Each predictor was evaluated in combination with all 15 gene feature sets. For transcriptomic-based approaches, we implemented a modular framework capable of integrating different gene features^59^ and evaluated it using the strongest gene feature identified in the direct prediction experiments. Details of the predictor implementations are provided in Note S1.

In addition to these methods, four approaches could not be summarized by the two-step framework described above. These include two network simulation–based methods (for which gene features alone do not fully characterize the approach), a method that directly assigns proportions from the training perturbation with the closest Gene Ontology annotation (and therefore does not use an explicit predictor), and a method that uses cells with the target gene lowly expressed to predict the cell state proportion. Detailed descriptions of their implementations are provided in Note S1.

### Benchmark on prediction error (Challenge 1)

For a robust comparison between the different methods, we chose a cross-fold validation setup instead of relying solely on the seven held-out perturbations in Challenge 1. In this setup, we randomly partitioned the perturbations in each experiment into *k* folds, and then ran each method by holding out each of the *k* fold, and training on the remaining folds, and testing on the held-out fold. The final accuracy of each method is evaluated based on the averaged test result over *k* folds, where we repeat this procedure 5 times to evaluate the standard deviation of the accuracy.

The number of folds *k* depends on the number of perturbations of the experiment; we set it to 6 for Screen 1 (with a total of 59 perturbations that passed preprocessing) and the public screen (with a total of 173 perturbations that passed preprocessing), and 5 for Screen 2 (with a total of 50 perturbations that passed preprocessing). We considered the absolute error as the measure for accuracy, as defined above in Challenge setup and results. The hyperparameters for each method and how they were selected for each run are provided in Note S1.

### Benchmark on ranking genes (Challenge 2)

To benchmark different methods for Challenge 2, we followed a similar procedure. In particular, for each method, we used all perturbations in Screen 1 as training data to predict the cell state proportion for the perturbations tested in Screen 2. These predictions were then used to rank the perturbations using the two objective functions, *f*_*ICB*_ and *f*_*CAR-T*_, and compared to the ground-truth rankings based on experimental data in Screen 2 to compute the AUC score, as defined above in Challenge setup and results. Each method was run 5 times to evaluate the standard deviation of the AUC score. The hyperparameters for each method and how they were selected for each run are provided in Note S1.

### Feature analyses

For computing the canonical correlation coefficients in Figure 2B and Figure S3B, we used PLSCanonical^70^ with one component. For the scBERT, Gene2Vec, GeneVector, and GO-node2vec features, as their dimensions far exceed the dimensions of other features, we reduced their dimensions to the top 10 principal components to avoid overestimation in the resulting canonical correlations. In the analysis of GWPS_K562 features in Figure S3A, we computed gene similarities in each dataset by computing the Pearson correlation between the pseudo-bulk expressions of each pair of perturbations. In the analysis of exon features in Figure S3C-E, the log-fold change for each perturbation was computed based on the averaged expressions of the target gene both under the corresponding perturbation and in the control cells. The log-expression of each gene was computed by averaging all the control cells. We predicted log-fold changes or log expressions using linear regression, where a random 20% of the genes were held-out as the test set. In Figure S4C, we computed log expressions by averaging all cells in the unperturbed PBMC dataset.^49^ In Figure S4D-E, we computed the Pearson correlations between each exon feature and the CRISPR gene effect in DepMap.^50^ In Figure S4F-G, we used the omics expressions of protein-coding genes in *log*(*TPM* + 1) format from DepMap.^50^

Since the exon features were mostly correlated with on-target effect sizes (*r* ≥ 0. 10, Figure S4A-B), we ran conditional independence tests on our dataset to test if the log-fold changes of target genes and their exon features were independent given the log expression of these genes in the control cells. For the conditional independence test, we used a nearest-neighbor estimator of conditional mutual information for discrete-continuous mixtures.^71,72^ We found that the variables were not conditionally dependent, with *p*_*val*_ = 0. 147, 1. 0, and 0. 963 for exon count, mean length, and maximum length, respectively.

### Analysis of the two hits

To search for similar genes in the pathway analysis for the two hits, we analyzed four different sources specified below. (1) We used the control cells in Screen 2 to identify co-expressed genes. In particular, for *Ndufv2* and *Dimt1*, we computed the absolute correlation coefficient between their expression and the expression of each expressed gene across all control cells. The top 50 genes with the highest absolute correlations were defined as the co-expressed genes for *Ndufv2* and *Dimt1*, respectively. (2) We obtained co-clustered genes in a genome-wide OPS screen on STING trafficking.^56^ For this, we downloaded the extracted features per perturbation and computed the absolute cosine similarity between the features of *Ndufv2* and *Dimt1* and the features of each perturbation in the dataset. The top 50 genes with the highest cosine correlations were defined as the co-clustered genes for *Ndufv2* and *Dimt1*, respectively. (3) We then obtained the dependency genes with *Ndufv2* and *Dimt1* in DepMap^50^ using the extracted CRISPR gene effects and the RNAi gene effects separately. In particular, for the extracted CRISPR and RNAi datasets, we computed the absolute correlation coefficient between the effects of knocking out *Ndufv2* and *Dimt1* and the effects of knocking out other genes across different cell lines. For the CRISPR and RNAi datasets separately, the top 50 genes with the highest correlation coefficients were defined as the dependency genes for *Ndufv2* and *Dimt1*, respectively. (4) Finally, we computed the co-clustered genes with these two hits in the genome-wide perturb-seq screen^48^. For this, we computed the pseudo-bulk expression of each perturbation and subtracted the pseudo-bulk expression of the control cells to obtain a gene-effect vector for each perturbation. We then computed the absolute cosine similarities between the gene-effect vectors of perturbing *Ndufv2* and *Dimt1* and the gene-effect vectors of perturbing other genes. The top 50 genes with highest absolute cosine similarities were defined as the co-clustered genes for *Ndufv2* and *Dimt1*, respectively.

For *Ndufv2* and *Dimt1* respectively, the genes that appeared from at least two sources defined above were collected, where the CRISPR gene effects and RNAi gene effects from DepMap^50^ were considered as two sources. For each hit, we then extracted a protein-protein interaction network from STRING^30,31^ between these collected genes and the corresponding hit by setting the species to mouse and the minimum score threshold to 400. We then took the connected genes found in the network and identified their common downstream genes in the genome-wide Perturb-seq screen^48^. In particular, for the connected genes found in the network that were perturbed in the genome-wide perturb-seq screen^48^, we computed the differentially expressed genes by comparing cells under any of these perturbations against the control cells.

### Statistical analyses

Throughout the manuscript, we made use of the following statistical tests^73^: one-sided Mann-Whitney U tests in Figure S1G (1036 coefficients for the same target gene, and 53,720 coefficients for different target genes), Figure S5A-B (33 shared targets), Figure 6C (19,890 all genes, 120 overlapping genes); Pearson correlation test in Figure S1H (74 targets); Spearman correlation test in Figure 3F (15 methods), and Figure 6B (120 overlapping genes); t-test with overestimated variance in Figure 6A (only control cells were considered: total of 4503 cells, 1957 cycling cells, 628 effector cells). The test results are specified in the text and figure legends.

