## Supplementary material for "A community machine learning challenge to predict the effects of gene perturbations on T cell differentiation for cancer immunotherapy": Note S1. Implementation of methods and hyperparameter selection.

### Supplementary Note S1 for A community machine learning challenge to predict the effects of gene perturbations on T cell differentiation for cancer immunotherapy

Author: Jiaqi Zhang[1,2], Marc A Schwartz[3], Mohammed Mutaher[3], Oluwatomisin Olajide[3], Yuri Pritykin[4], Orr Ashenberg[1,5]*, Nir Hacohen[3,6,]*, Caroline Uhler[1,2,]*

[1]Eric and Wendy Schmidt Center, Broad Institute of MIT and Harvard, Cambridge, 02142, MA, USA.
[2]Laboratory for Information and Decision Systems, Massachusetts Institute of Technology, Cambridge, 02139, MA, USA.
[3]Broad Institute of MIT and Harvard, Cambridge, MA, 02142 USA
[4]Department of Computer Science and Lewis-Sigler Institute for Integrative Genomics, Princeton University, Princeton, NJ 08540, USA
[5]Klarman Cell Observatory, Broad Institute of MIT and Harvard, Cambridge, 02142, MA, USA
[6]Krantz Family Center for Cancer Research, Massachusetts General Hospital, Charlestown, 02129, MA, USA

#### Implementation of gene features derived from top submissions

Based on the submitted writeups and code (Table S2), we reimplemented the following gene features, with renamed labels summarizing each approach, detailed below.

**Expression_Conditioned**. This approach relies on the hypothesis that genes that are co-expressed across different conditions will exhibit similar perturbational effects when perturbed. It represents each gene by its expression across multiple conditions in the training data, including unperturbed and perturbed conditions. In particular, suppose that in the training set we have access to unperturbed cells $C^{NT}$ and perturbed cells $C^1, ..., C^k$ corresponding to $k$ different perturbations. For a given gene $i$, whose expression is measured in each cell, the **Expression_Conditioned** feature represents $i$ as a $(k+1)$-dimensional vector $R_i$. The first entry of $R_i$ is the mean expression of gene $i$ across $C^{NT}$, and the subsequent $k$ entries are the mean expressions of gene $i$ across $C^1, ..., C^k$, respectively. This construction provides meaningful feature representations for all expressed genes; genes not measured in expression are assigned an all-zero $(k+1)$-dimensional vector.

**5Features.** This approach is similar to **Expression_Conditioned**, but instead of focusing on co-expression across different perturbation conditions, it emphasizes expression patterns across cell states. The participants proposed to select features from summary statistics of gene expression, together with the expression of cell state marker genes: *Tcf7*, *Bach2*, *Prf1*, *Gzma*, *Pdcd1*, and *Eomes*. They ended up with a total of five features. Formally, let the training cells be denoted by $C$, with "progenitor", "effector", "cycling", and "other" subsets $C^{prog}$, $C^{eff}$, $C^{cyc}$, and $C^{other}$, respectively. For a given gene $i$, whose expression is measured in each cell, **5Features** represents gene $i$ by a 5-dimensional vector $R_i$. The first entry of $R_i$ is the standard deviation of the sum of expression of gene $i$ and *Tcf7* across $C$, and the remaining four entries are the median expression of gene i across $C^{prog}$, $C^{eff}$, $C^{cyc}$, and $C^{other}$, respectively. Same as above, genes that are not measured in expression are assigned an all-zero 5-dimensional vector.

**14Features.** This approach is similar to **Expression_Conditioned**, but instead of computing a single statistic for each individual perturbation, it computes multiple statistics aggregated across all perturbations. In particular, denote in the training set the unperturbed cells as $C^{NT}$ and perturbed cells as $C^{Pert}$. For a given gene $i$, whose expression is measured in each cell, **14Features** represents it by a 14-dimensional vector $R_i$, constructed as the concatenation of seven statistics computed on $C^{NT}$ and the same seven statistics computed on $C^{Pert}$. The seven statistics for $C^{NT}$ (and similar for $C^{Pert}$) are: the ratio of cells with nonzero expression of gene $i$ in $C^{NT}$ ($C^{Pert}$); the mean, standard deviation, and skewness of gene $i$'s expression across $C^{NT}$ ($C^{Pert}$); and the mean, standard deviation, and skewness of gene $i$'s expression restricted to cells with nonzero expression. As above, genes that are not measured in expression are assigned an all-zero 14-dimensional vector.

**Matrix_Completion.** This approach constructs gene features in three steps. Denote the cell-by-gene expression matrix for training as $M \in \mathbb{R}^{N\times G}$, where $N$ is the number of cells and $G$ is the number of expressed genes. In the first step, the method identifies a subset of expressed genes whose expression is correlated with cell number or cell state distribution, denoted by $S$. In particular, for each expressed gene, a $(k + 1)$-dimensional vector is constructed that records its mean expression across unperturbed cells and across each of the $k$ perturbations in the training set. Similarly, a $(k + 1)$-dimensional vector is constructed that records either the cell counts or the proportions of the five cell states across unperturbed cells and the $k$ perturbations. The Pearson correlation between these two vectors is then computed, and a gene is included in $S$ if and only if the correlation exceeds $0.5$.

In the second step, the correlation between each expressed gene and each gene in $S$ is computed across cells in $M$, resulting in a matrix $D \in \mathbb{R}^{G\times|S|}$. This matrix is then projected onto its first $100$ principal components, yielding $D^{PC} \in \mathbb{R}^{G\times 100}$.

In the third step, the method learns transformation matrices $W, H \in \mathbb{R}^{100\times r}$ by minimizing the Frobenius norm between two matrices: (1) $[D^{PC}]_{k'} W H^{\top} [D^{PC}]_{k'}^{\top} \in \mathbb{R}^{k'\times k'}$ where $k'$ denotes the number of target genes amongst the $k$ training perturbations that are expressed, and $[D^{PC}]_{k'} \in \mathbb{R}^{k'\times 100}$ denotes the rows of $D^{PC}$ that corresponds to training perturbations whose target genes are expressed, and (2) $P \in \mathbb{R}^{k' \times k'}$. Each row of $P$ is a binary vector with five ones, indicating, for each of the $k'$ perturbation, the five perturbations whose cell state proportion vectors are most similar in $l_2$ norm. The rank parameter $r$ is set to $100$ to control the rank of $W, H$. The optimization is performed using conjugate gradient descent. After learning $W, H$, the method computes $D^{PC} W H^{\top} \in \mathbb{R}^{G \times 100}$ and uses the corresponding row as the feature representation for each expressed gene. Genes not measured in expression are assigned an all-zero $100$-dimensional vector.

**scETM.** This approach obtains gene embeddings by applying scETM[1] to the provided training data. scETM is an unsupervised topic-modeling method trained to reconstruct single-cell transcriptomic data. In the learned embedding space, each expressed gene is represented by a gene embedding of a chosen dimensionality, which was set to $32$ here, as the participants observed no significant drop in negative log-likelihood when increasing the dimension. After pretraining scETM on the training data, gene embeddings were extracted from the model parameters. Genes that were not measured in expression were assigned an all-zero $32$-dimensional vector. Finally, all gene embeddings were standardized by subtracting the mean and scaling to unit variance.

**scVI.** This approach is similar to **scETM**, but instead of using the topic model scETM, it uses the linearly decoded variational autoencoder (LDVAE)[2] implemented in scVI tools.[3] The latent dimensionality is set to $50$. After pretraining, gene embeddings are extracted from the loadings of the linear decoder. Genes that were not measured in expression were assigned an all-zero $50$-dimensional vector.

**PPI_Weighted.** This approach consists of two steps. In the first step, an embedding is constructed for each condition in the training set. Denote the unperturbed cells by $C^{NT}$ and the perturbed cells in the training set by $C^1, \ldots, C^k$, corresponding to $k$ different perturbations. The method first identifies genes that are differentially expressed across the five cell states using Scanpy's implementation with default parameters;4 these genes are denoted by $DE$. For each condition, it then computes four $DE$-dimensional vectors, denoted by $(R^{NT,prog}, R^{NT,eff}, R^{NT,cyc}, R^{NT,term})$ for the unperturbed condition and $(R^{l,prog}, R^{l,eff}, R^{l,cyc}, R^{l,term})$ for $l = 1, \ldots, k$. Each vector is obtained by selecting cells from the corresponding condition and one of the four states—progenitor, effector, cycling, and terminally exhausted—and averaging the expression of genes in $DE$. An embedding is then computed for each condition, denoted by $R^{NT}, R^1, \ldots, R^k$. For the unperturbed condition, $R^{NT}$ is obtained by averaging $(R^{NT,prog}, R^{NT,eff}, R^{NT,cyc}, R^{NT,term})$. For $l = 1, \ldots, k$, $R^l$ is obtained by a weighted average of $(R^{l,prog}, R^{l,eff}, R^{l,cyc}, R^{l,term})$, where the weights are given by the average expression of the target gene among cells in $C^l$ of the corresponding state when the target gene is measured in expression; otherwise, equal weights of $1$ are used.

In the second step, for each gene $i$, **PPI_Weighted** identifies training perturbations whose target genes are associated with i using the STRING Mus musculus protein interaction network,[4] denoted by $S_i$. This is determined using a multi-step criterion: if there exist target genes with a combined STRING score greater than $600$, then $S_i$ consists of all such genes; otherwise, if there exist target genes with a combined score greater than $0$, $S_i$ consists of the target genes with the highest scores; otherwise, $S_i$ is empty. Finally, the feature representation $R_i$ for gene $i$ is computed as the average of $R^l$ over all $l$ in $S_i$ if $S_i$ is nonempty; otherwise, $R_i$ is set to $R^{NT}$.

**GWPS_K562_H2M.** This approach utilizes the genome-wide Perturb-seq screen (GWPS),[5] which measures the effects of 9,866 gene knockdowns in human K562 cells. It builds a mapping from human gene knockdown effects to their mouse ortholog gene knockout effects using the training data. In particular, for each knockdown targeting gene $I$ in GWPS, it computes the mean of the first 50 principal components across all cells with knockdown $I$, denoted as $R_I^{human}$. As a special case, it also computes $R_0^{human}$ for unperturbed cells. Similarly, it computes $R_i^{mouse}$ for each training perturbation $i = 1,..., k$, as well as for the unperturbed cells $i = 0$.

An MLP regressor with a single hidden layer of latent dimension $10$ is then trained to map from $R_I^{human}$ to $R_i^{mouse}$ for $i = 0$ and for all $i = 1,..., k$ such that the corresponding human ortholog $I$ is perturbed in GWPS. Finally, the trained MLP is applied to $R_I^{human}$ for $i = 0$ and for all genes perturbed in GWPS to obtain the corresponding features $R_i$, where $i$ denotes the mouse ortholog. Genes whose features cannot be obtained through this process, either because no human ortholog exists or because the human ortholog was not perturbed in GWPS, are assigned the unperturbed representation $R_0$.

**GWPS_K562.** This approach is similar to **GWPS_K562_H2M,** but it did not use the training data, and instead directly use $R_I^{human}$ as the gene feature $R_i$ for the mouse ortholog $i$ of $I$. As above, genes whose features cannot be obtained through this are assigned the unperturbed representation $R_0^{human}$.

**Exons.** This approach uses exon annotation information downloaded from NCBI[6] to construct gene features. For each gene $i$, the feature is a $6$-dimensional vector consisting of summary statistics of exon lengths within the gene region: the minimum, maximum, mean, median, first quartile, and third quartile of the exon lengths.

**scBERT.** This approach extracts pretrained human gene embeddings from scBERT [7] and uses it directly to represent their mouse orthologs, where each gene feature is a $100$-dimensional vector. Genes whose features cannot be obtained through this are assigned all-zero $100$-dimensional vector.

**Gene2Vec.** Similar as above, this approach directly uses pretrained human gene embeddings, but instead from the gene2vec model,[8] where each gene feature is a $200$-dimensional vector. Genes whose features cannot be obtained through this are assigned all-zero $200$-dimensional vector.

**GeneVector.** Similar as above, this approach directly uses pretrained human gene embeddings, but instead from the GeneVector model,[9] where each gene feature is a $128$-dimensional vector. Genes whose features cannot be obtained through this are assigned all-zero $128$-dimensional vector.

**GO-node2vec.** This approach obtains gene features by training a graph neural network with the gene-ontology database. It first generates an undirected, unweighted graph with genes as nodes and edges representing genes that share the same pathway. It then trains Node2Vec implemented in Pytorch Geometric,[10] to learn a $128$-dimensional embedding for each gene. Genes whose features cannot be obtained through this are assigned all-zero $128$-dimensional vector.

#### Implementation of our proposed gene features

Our proposed **Perturbed_Tcell** gene features are constructed by concatenating a selective subset of individual features after minimax normalization each individual feature from the following sources: proliferation scores under various conditions,[11–15] production scores of various markers,[16,17] time-coarse bulk differential expression,[18] and base-editing screen with clinical annotations.[19]

There were a total of 28 gene features available for selection, which are described in detail in the next paragraph. Among these 28 features, we performed feature selection using a greedy approach on the cross-validation splits described in the “Benchmark on Challenge 1” section of Methods. This greedy procedure starts with all 28 features and successively removes one existing feature at a time, selecting the feature whose removal yields the largest decrease in the average cross-validation loss and exceeds a pre-set tolerance (0.005). When no further feature can be removed under this criterion, the procedure then successively adds one feature not currently included, choosing the feature whose addition yields the largest decrease in the average cross-validation loss and exceeds the same tolerance. The algorithm alternates between removal and addition steps for a maximum of 10 rounds, where one round consists of a full removal phase followed by a full addition phase. This process resulted in the selection of 24 gene features, and therefore not many features were excluded.

We now provide details on how gene features were extracted from each dataset and whether they were retained for the final feature concatenation. For all datasets, genes with missing features were imputed using the average of the available gene features. For the proliferation screen,[11] we included the log2 ratios from the metabolic and Brie libraries, resulting in two scalar features for genes included in each library. We excluded the feature derived from the metabolic library. For the time-coarse bulk differential expression dataset,[18] we computed bulk differential expression for all pairs of conditions (naive, day 4, and day 7 under chronic or acute infection), except for chronic day 7 versus acute day 7, resulting in nine scalar features per measured gene. We excluded the feature corresponding to chronic day 7 versus chronic day 4. For the proliferation screen,[12] we used the log fold change from the genome-wide library, yielding one scalar feature per gene. We additionally included all log fold changes from the minipool library, resulting in a 7-dimensional feature per gene. For the production screen,[16] we extracted the control mean, treatment mean, and log fold change for each of four markers (*TNFα*, *CD25*, *IFNγ*, and *PD1*), resulting in four 3-dimensional feature blocks per gene. For the proliferation screen,[13] we used all seven measured statistics from the ORF screen, yielding a 7-dimensional feature per gene. For the proliferation screen,[14] we used the negative log fold changes under four treatments—Adenosine, Cyclosporine, Tacrolimus, and TGFβ—resulting in four scalar features per gene. We excluded the feature corresponding to Tacrolimus. For the base-editing screen,[19] we included 26 clinically annotated statistics per gene, resulting in a 26-dimensional feature per gene. For the proliferation screen,[15] we used all three log fold changes per gene from the metabolic library, resulting in a 3-dimensional feature per gene; this feature set was excluded from the final concatenation. Finally, for the production screen,[17] we included gene effect scores from CRISPRa and CRISPRi screens, as well as the negative log fold change from CRISPR knockout screens with and without additional perturbation of *BATF3*.

#### Implementation of predictors

The 2-step methods first find a gene feature, described in the last section, and then construct a predictor that maps gene feature to cell state proportion. For the predictor in the second step, we considered two types of approaches, direct and transcriptomic-based.

For the direct approaches, we included three predictors, each tested in combination with all fifteen gene features described above, resulting in a totality of fourty-five methods, presented in the main text. The three predictors are: (1) A nearest neighbor regressor with uniform weights and a neighborhood size $k \in \{2, ..., 30\}$, where $k$ is selected based on cross-validation described in the section on hyperparameter selection. This method predicts the cell state proportion of a held-out perturbation based on averaging the cell state proportions of the nearest $k$ perturbations to the held-out one in terms of distances between their gene features. The minkovski distance is used if the gene features are of dimension $\leq 2$, otherwise the correlation based distance is used. The gene features are optionally reduced to their first 50 principal components before the distance is computed, depending on the cross-validation loss described below. (2) A kernel ridge regressor is used with $\{laplacian, linear, rbf, cosine\}$ kernel and $l_2$ penalty weight in $\{0.001, 0.01, 0.1, 1, 10\}$, where the choice of kernel and $l_2$ penalty weights are selected based on cross-validation described in the section on hyperparameter selection. The gene features are optionally reduced to their first 50 principal components or min-max standardized before the kernel regressor is applied, depending on the cross-validation loss described below. (3) A bagging regressor is used with $\{2, 4, 8\}$ kernel ridge estimators with the same kernel and $l_2$ penalty weight, which are chosen from the same set as described above. The number of estimators, the kernel, and the $l_2$ penalty weights are selected based on cross-validation described in the section on hyperparameter selection. The gene features are reduced to their first 50 principal components if they are of dimension $> 50$ and optionally min-max standardized before the kernel regressor is applied, depending on the cross-validation loss described below

For the transcriptomic-based approaches, we implemented two predictors, one using the **Perturbed_Tcell** feature and one using the **GWPS_K562_H2M** feature, resulting in two methods: (1) one uses the **Perturbed_Tcell** feature within MORPH,[20] a modular framework that uses gene features as input to predict the transcriptomic distributions of held-out perturbations. Here we consider predicting the expressions of top 1,000 highly variable genes. We then train an MLP classifier using the control cells to predict cell state from the expression of these 1,000 highly variable genes. This classifier is then applied to samples from the predicted transcriptomic distribution of a held-out perturbation to obtain predicted cell states of these samples. These predicted cell states are then used to compute the predicted cell state proportion of the held-out perturbation. (2) Another method uses the **GWPS_K562_H2M** feature, but instead of mapping the feature directly to state proportions, it first samples from a multi-variate gaussian distribution centered at the feature, which is the predicted principal components of perturbations, with variance being the variance of the principal components of control cells in the training data. Then similarly as in the first method, a classifier is trained to map the sampled PCs to cell states.

#### Implementation of additional methods

We additionally implemented four methods in Figure S2a. One of the four methods directly predicts cell state proportions; the remaining three methods are transcriptomics-based, where knock-out effects on transcriptomics are predicted and then classified into cell states.

In particular, **GO_knn** first constructs a similarity matrix between genes perturbed in the training set and genes perturbed in the held-out test set, where each entry represents the number of GO terms[21,22] in which the corresponding pair of genes both appear. It then predicts, for each held-out perturbation, the weighted average of the cell state proportions of its three most similar training perturbations, where the weights are given by the similarity matrix entries. For genes that are not found in any GO terms, the control cell state proportion is used as the prediction.

**LowExp** first predicts the transcriptomic response of a held-out perturbation if the corresponding gene has more than 1,000 cells with nonzero expression in the training data, using the 5,000 cells in the training data with the lowest expression of that gene. The predicted cell state proportions are then obtained from the state annotations of these 5,000 cells. If the corresponding gene has no more than 1,000 cells with nonzero expression in the training data, the control cell state proportions are used as the prediction.

**scTenifoldNet** first constructs a network based on control cells and performs a virtual gene knockout by setting the gene's outgoing edges to zero in the network, following.[23] To obtain the resulting proportions, it first selects the 128 genes that are most predictive of cell states using

SelectKBest.[24] It then trains a random forest classifier to map these 128 genes to cell states and obtains the corresponding feature importances. For each gene knockout $i$, it selects the 55 genes with the highest feature importances for each state $s$ and averages their fold changes predicted by the virtual gene knockout to obtain $c_i^s$. It then computes a min–max normalized score $d_i^s = (c_i^s - \min_{s'} c_i^{s'}) / (\max_{s'} c_i^{s'} - \min_{s'} c_i^{s'})$ for each state $s$, and predicts the proportion of state $s$ as $d_i^s$ divided by the sum of $d_i^{s'}$ over all possible $s'$. For genes for which no knockout can be simulated, the control cell state proportions are used as the prediction.

**Interv** first imputes the zero-inflated expression matrix using MAGIC.[25] It then uses the training data to classify cell states from gene expression using LASSO[26] and identifies a subset of genes $S$ with nonzero coefficients. To predict the effect of a knockout of gene $i$, it uses control cells to identify a Markov blanket $T$ of $i$, and then identifies the parents $pa(i)$ of $i$ from the Markov blanket using local LiNGAM.[27] For each gene $j$ in $S$, it regresses $j$ on $i \cup pa(i)$ using control cells to estimate the total causal effect from $i$ to $j$. It then computes regression residuals for genes in $S$ and subtracts $i$'s contribution from $S$ to obtain the predicted transcriptomic profile under knockout of gene $i$ in control cells. These predicted transcriptomic profiles are then mapped to cell states using the LASSO classifier, and the corresponding proportions are computed. For genes that are not expressed, the control cell state proportions are used as the prediction.

#### Hyperparameter selection

We select the hyperparameters of the predictors based on a cross-fold validation scheme on the training data. This applies to both the train–test splits in Figure S2b for challenge 1, detailed in the "Benchmark on prediction error (challenge 1)" section of Methods, and the benchmark for challenge 2, detailed in the "Benchmark on prediction error (challenge 1)" section of Methods. For both cases, we randomly partition the training data into five folds. For a set of hyperparameters, we perform five-fold cross-validation by leaving out one fold, training on the remaining folds using this set of hyperparameters, and computing the accuracy on the left-out fold. The final score of this set of hyperparameters is the averaged accuracy across the five folds. We then select the set of hyperparameters with the highest score.
